# Excessive Polyamine Generation in Keratinocytes Promotes Self-RNA Endosomal Sensing by Dendritic Cells in Psoriasis

**DOI:** 10.1101/2020.03.09.984658

**Authors:** Fangzhou Lou, Yang Sun, Zhenyao Xu, Liman Niu, Zhikai Wang, Siyu Deng, Zhaoyuan Liu, Hong Zhou, Jing Bai, Qianqian Yin, Xiaojie Cai, Libo Sun, Hong Wang, Zhouwei Wu, Xiang Chen, Yuling Shi, Wufan Tao, Florent Ginhoux, Honglin Wang

**Author notes:** Correspondence to: Honglin WANG, Ph.D. Shanghai Institute of Immunology, Shanghai Jiao Tong University School of Medicine (SJTU-SM), 280 South Chongqing Road, 200025 Shanghai, China, Florent GINHOUX, Ph.D., Singapore Immunology Network, Agency for Science, Technology and Research, 8A Biomedical Grove Level 3, Immunos Building, Singapore 138648, Singapore. These authors contributed equally (Fangzhou Lou, Yang Sun).

## Abstract

The mechanisms underlying tissue-specific chronic inflammation are elusive. Here we report that mice lacking Protein Phosphatase 6 in keratinocytes are predisposed to psoriasis-like skin inflammation, with an inordinate urea cycle and enhanced oxidative phosphorylation that supports hyperproliferation. This phenotype is mediated by increased Arginase-1 production resulting from CCAAT/enhancer-binding protein beta activation. Single-cell RNA-seq of the psoriatic epidermis revealed that the rate-limiting enzyme for Arginine biosynthesis, Argininosuccinate synthetase 1, maintains the Arginine pool, which is indispensable for immune responses. Accumulated polyamines branched from the urea cycle promote endosomal Tlr7-dependent self-RNA sensing by myeloid dendritic cells. This process is achieved with the assistance of an RNA-binding peptide that originates from the heterogeneous nuclear ribonucleoprotein A1, a probable autoantigen in psoriasis. Finally, targeting urea cycle wiring with an arginase inhibitor markedly improved skin inflammation in murine and non-human primate models of psoriasis. Our findings suggest that urea cycle alteration and excessive polyamine production by psoriatic keratinocytes promote self-RNA sensing by dendritic cells, which links the hyperproliferation of stationary cells with innate-immune activation in an auto-inflammatory condition.

## INTRODUCTION

The regulation of immune cell-intrinsic metabolism, or “immunometabolism”, is under extensive investigation. The goal of such research is to identify translational applications of manipulating metabolism to treat immune-mediated diseases. One example is the targeting of oxidative phosphorylation (OXPHOS) to impair T-helper 17 (Th17) cell generation *in vivo* in a mouse model of psoriasis; here, the endogenous metabolite itaconate inhibits the IL-17-IκBζ axis and ameliorates psoriasis-like skin inflammation (Bambouskova et al., 2018; Franchi et al., 2017). Although these therapeutic approaches are effective, psoriasis always relapses and is still incurable in human patients due to our incomplete understanding of the causes of the type 17-biased immune responses in the skin.

As a skin-confined chronic inflammatory condition, psoriasis is characterized by the abnormal interplay between hyperproliferative epidermal keratinocytes and self-reactive immune cells (Lowes et al., 2007). Although psoriasis has long been regarded as a metabolism-associated disorder (Haslam and James, 2005), and is assumed to be fueled by keratinocyte-intrinsic metabolic fluxes, the choreography of metabolic adaptions in psoriasis and the underlying mechanisms are poorly defined. Abnormal amino acid and lipid metabolism was noticed in patients with psoriasis (Gudjonsson et al., 2009; Kamleh et al., 2015; Kang et al., 2017), with dietary saturated fatty acids shown to exacerbate the skin inflammation (Herbert et al., 2018).

Meanwhile, inhibiting glucose transportation in keratinocytes seems to be a novel means for treating psoriasis (Zhang et al., 2018). Whether there exists a tissue-specific metabolic scenario and if so, what the functional significance in the context of inflammation might be, however, is unknown.

The other aspect of the auto-inflammatory loop in psoriasis pathogenesis is innate immunity activation, which triggers subsequent adaptive immune responses against self-components. LL-37, ADAMTSL5, PLA2G4D and HNRNPA1 are probable psoriasis autoantigens (Arakawa et al., 2015; Cheung et al., 2016; Guarneri et al., 2018; Jones et al., 2004; Lande et al., 2014), with LL-37 and HNRNPA1 harboring RNA-binding properties. Given the notion that self-RNA sensing is central to the development of inflammatory diseases like psoriasis and lupus (Celhar et al., 2015; Ganguly et al., 2009), understanding how these RNA-binding proteins are handled by antigen-presenting cells before they become recognized by self-reactive T cells should be addressed.

We previously reported that a microRNA directly targets Protein Phosphatase 6 (PP6) as a negative cell cycle regulator in psoriatic keratinocytes (Yan et al., 2015). PP6, which consists of one catalytic subunit and two regulatory subunits, is crucial for skin homeostasis (Bonilla et al., 2016; Hodis et al., 2012; Krauthammer et al., 2012). Given the versatility of serine/threonine phosphatases, we aimed to determine the specific functionality of PP6 in keratinocytes. Using a genetic ablation approach, we discovered that PP6 functions as an essential check point in epidermal metabolic regulation and in psoriasis development. Our delineation of the transcriptional and metabolic rewiring that occurs subsequent to PP6 deficiency might help illuminate the metabolite-facilitated activation of innate immunity in psoriasis and other auto-inflammatory disorders.

## RESULTS

### Epidermis-specific Pp6-deficient mice spontaneously develop psoriasis-like skin inflammation

We previously found that PP6 is directly targeted by microRNA-31, one of the most highly overexpressed microRNAs in human psoriatic skin (Xu et al., 2013; Yan et al., 2015). We first performed an immunohistochemical analysis of PP6 on skin sections derived from five healthy donors and 117 patients with psoriasis. We found that PP6 was ubiquitously expressed in all epidermal layers of the healthy skin but was comparatively reduced in psoriatic lesions (**Figure S1A**). Notably, we detected a strong negative correlation between PP6 levels and epidermal hyperplasia (**Figure S1B**). When correlating the Psoriasis Area and Severity Index (PASI) and the PP6 expression levels, we found that epidermal PP6 levels in the severe group (PASI > 20) were significantly lower than the levels in the mild and moderate groups (*p* < 0.05, **Figure S1C**). We also found decreased PP6 expression in both imiquimod (IMQ)-induced and IL-23-induced mouse models of psoriasis (**Figures S1D-S1G**).

We next crossed mice with loxP-flanked *Pp6* alleles (Pp6^fl/fl^) with Keratin 5-Cre (K5) mice to specifically delete *Pp6* in keratinocytes (designated K5. Pp6^fl/fl^, **Figures S2A-S2C**). K5^+^ medullary thymic epithelial cells (mTECs) did not express Pp6 protein and were likely unaffected by *Pp6* genetic ablation (**Figure S2D**). K5. Pp6^fl/fl^ mice were viable, and mice younger than 16-week old exhibited no notable skin abnormalities (data not shown). Strikingly, skin lesions spontaneously emerged in K5. Pp6^fl/fl^ mice after 16 weeks of age (**Figure 1A**). We found scaly plaques on the faces, ears, upper backs and tails of the affected mice (**Figure 1B**). Histological examination of the skin lesions showed remarkable epidermal hyperplasia (acanthosis) with loss of the granular layer, hyperkeratosis and parakeratosis in the epidermis together with microabscesses that accumulated on the surface of the thickened epidermis and massive cellular infiltrates in the dermis (**Figures 1C** and **1D**). Splenomegaly and lymphadenopathy were also pronounced in the affected K5. Pp6^fl/fl^ mice (**Figure S2E**).

**Figure 1.**
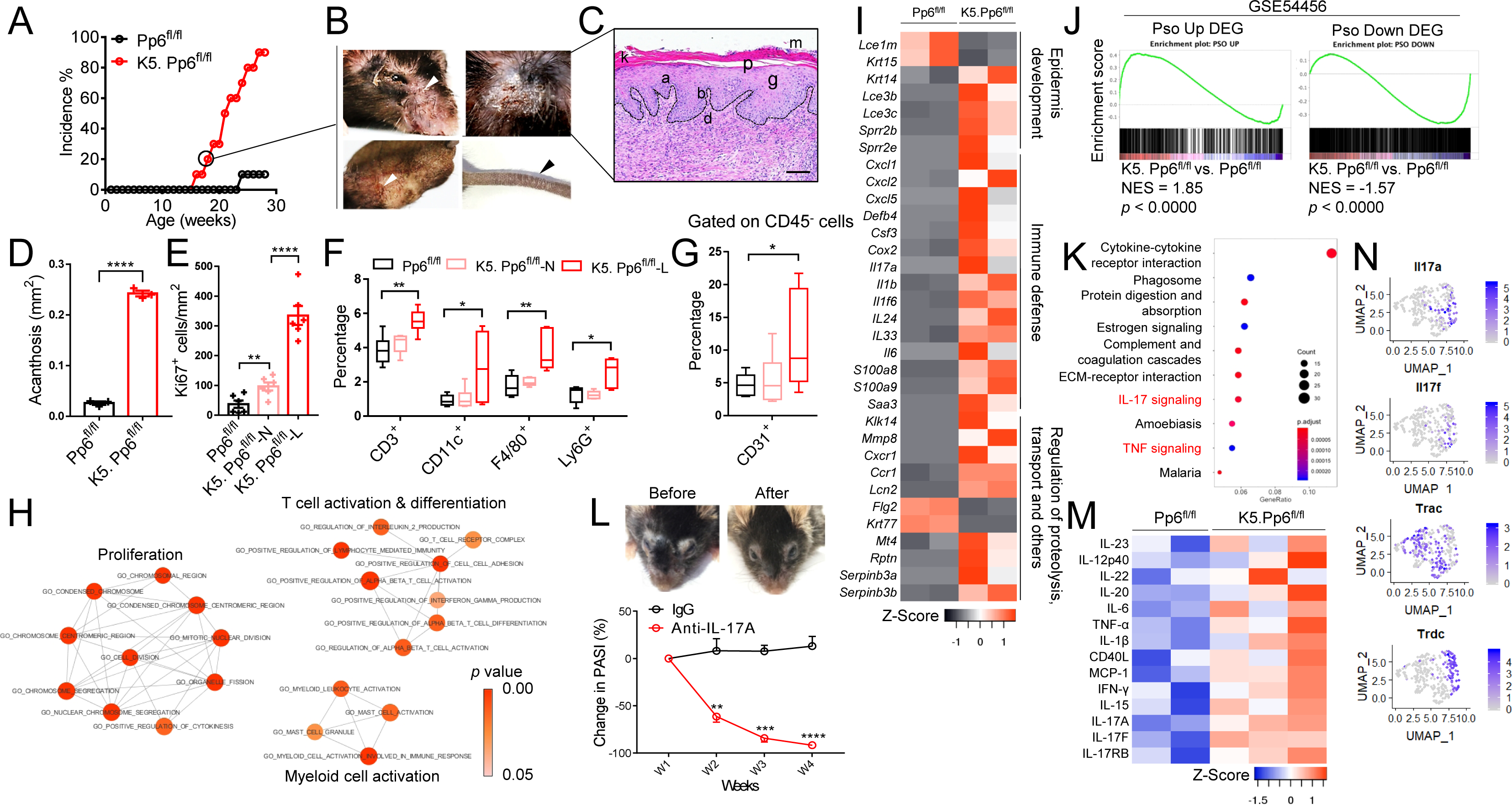
Mice lacking Pp6 in the epidermis show hallmarks of psoriasis. (**A**) The incidence of skin lesions in K5. Pp6^fl/fl^ mice and Pp6^fl/fl^ mice over time (n = 10). (**B**) Macroscopic views of the face, ear, upper back and tail skin from an 18-week-old K5. Pp6^fl/fl^ mouse. Scaly plaques are marked by arrowheads. (**C**) H&E staining of the skin lesions from the upper back of a K5. Pp6^fl/fl^ mouse. a, acanthosis; b, increased proliferative basal layer epidermal keratinocytes; d, dermal cell infiltrates; k, hyperkeratosis; p, parakeratosis; g, loss of stratum granulosum; m, Munro’s microabscess. The dotted line indicates the border between the epidermis and the dermis; scale bar represents 100 μm. (**D**) Acanthosis of diseased K5. Pp6^fl/fl^ skin compared to Pp6^fl/fl^ skin (n = 4). (**E**) Quantitation of Ki67^+^ epidermal cells in mouse skin sections (n = 6). (**F**) Statistics of the flow cytometric analysis of immune cell-specific markers in mouse skin (n = 5∼6). The boxes show the 25th to 75th percentiles; the whiskers show the minimum to maximum; and the lines show the medians. (**G**) Flow cytometric analysis of CD31^+^ cell percentages in the CD45^-^ cell compartments in mouse skin (n = 5∼6). The boxes show the 25th to 75th percentiles; the whiskers show the minimum to maximum; and the lines show the medians. (**H**) GSEA of Gene ontology (GO) pathways identified by scRNAseq of Pp6-deficient keratinocytes compared to WT keratinocytes. *p* value Cutoff = 0.005; FDR Q-value Cutoff = 0.1. Networks with > 3 nodes are shown. (**I**) Heat map of selected genes based on RNA-seq data from the skin of K5. Pp6^fl/fl^ and Pp6^fl/fl^ mice (*p* < 0.05, Log2FC > 1.2, n = 2). The GO categories are indicated. (**J**) GSEA of DEGs in K5. Pp6^fl/fl^ skin relative to Pp6^fl/fl^ skin with upregulated (Pso Up) or downregulated (Pso Down) DEGs in human psoriatic skin (GSE54456) enriched. (**K**) Kyoto Encyclopedia of Genes and Genomes (KEGG) pathway enrichment of upregulated DEGs in K5. Pp6^fl/fl^ skin relative to Pp6^fl/fl^ skin. *p* value cutoff = 0.01; gene numbers enriched in each pathway are indicated by the sizes of the circles; the adjusted *p* values are indicated by the continuous color. (**L**) Macroscopic views of skin lesions in a representative K5. Pp6^fl/fl^ mouse before or after anti-IL-17A treatment (upper panel). Changes in PASI of diseased K5. Pp6^fl/fl^ mice on indicated time-point after anti-IL-17A treatment (lower panel). (**M**) IL-23-T17-associated cytokine levels detected by semiquantitative cytokine array of K5. Pp6^fl/fl^ and Pp6^fl/fl^ mouse skin. (**N**) Feature-plots of Il17a, Il17f, Trac and Trdc expression in T cells derived from K5. Pp6^fl/fl^ skin analyzed by scRNAseq. The data (**B** and **C**) are representative of > 8 mice. The data (**L**) are representative of two independent experiments with two mice in each group. Pp6^fl/fl^, healthy skin sample from control mice; K5. Pp6^fl/fl^-N, unaffected skin from K5. Pp6^fl/fl^ mice; K5. Pp6^fl/fl^-L, affected skin from K5. Pp6^fl/fl^ mice. \**p* < 0.05, \*\**p* < 0.01, \*\*\**p* < 0.001, \*\*\*\**p* < 0.0001, two-tailed Student’s *t*-test (means ± SEM).

We next characterized how skin lesions were initiated in K5. Pp6^fl/fl^ mice. We investigated barrier function [indicated by transepidermal water loss (TEWL)], keratinocyte proliferation (indicated by Ki67^+^ cell numbers in epidermis), Keratin 6 (K6) expression and CD3^+^ T-cell, CD11c^+^ dendritic cell (DC), F4/80^+^ macrophage and Ly6G^+^ neutrophil infiltration in non-lesional skin from Pp6^fl/fl^ mice, young K5. Pp6^fl/fl^ mice (10∼15-week age) and old K5. Pp6^fl/fl^ mice (≥ 16-week age), and lesional skin from old K5. Pp6^fl/fl^ mice (≥ 16-week age). We found that the number of Ki67^+^ cells was increased in the non-lesional epidermis from K5. Pp6^fl/fl^ mice (both young and old) compared to those of Pp6^fl/fl^, and in the lesional epidermis from K5. Pp6^fl/fl^ mice compared to the non-lesional epidermis from the same animals (**Figures S2F** and **1E**). Immune infiltration, TEWL and K6 expression were unaltered in the non-lesional skin from K5. Pp6^fl/fl^ mice (either young or old) but all parameters were increased in the lesional skin from K5. Pp6^fl/fl^ mice compared to Pp6^fl/fl^ controls (**Figures S2G, 1F**, **S2H** and **S2I**). The number of CD80^+^ activated antigen presenting cells was also increased in the non-lesional and lesional dermis of K5. Pp6^fl/fl^ mice (**Figure S2J** and **S2K**) and vascularization (indicated by CD31^+^ cell percentages) was increased in the skin lesions of K5. Pp6^fl/fl^ mice (**Figure 1G**). We found that the dermis of K5. Pp6^fl/fl^ mice was enriched in mast cells but lacked eosinophils; the serum IgE levels, however, were consistent with those of Pp6^fl/fl^ mice, suggesting a non-allergic reaction (**Figures S2L-S5N**).

We next performed single-cell RNA sequencing (scRNAseq) on epidermal cells derived from the ear skin of young K5. Pp6^fl/fl^ mice and C57BL/6 control mice, and compared the transcriptional profiles of the keratinocyte compartments with and without Pp6 deficiency. Using gene set enrichment analysis (GSEA) of gene ontology (GO) pathways and subsequent enrichment map visualization, we found that Pp6-deficient keratinocytes were highly proliferative and more involved in myeloid cell and T-cell activation processes, as compared with Pp6-intact keratinocytes (**Figure 1H**). These findings suggest that Pp6-deficient keratinocytes transduce proinflammatory signals to immune cells. Persistent environmental challenges to K5. Pp6^fl/fl^ mice might amplify the resulting inflammatory circuits to a certain threshold and underlie skin lesion development.

We next performed transcriptome profiling to study the altered genes in the K5. Pp6^fl/fl^ lesions. Among 18,835 changed genes, 1,619 were differentially expressed genes (DEGs; |log2FC| ≥ 1 and *p* < 0.05). We identified a notable resemblance between the gene expression patterns in the affected skin from K5. Pp6^fl/fl^ mice and the affected skin from patients with psoriasis, with similar expression profiles of psoriasis “countermark” genes. For example, keratinocyte-derived antimicrobial peptide (AMP) coding genes (*S100a7a*, *S100a8*, *S100a9*, *Lcn2* and *Defb4*) were strongly induced in the inflamed mouse skin (Gudjonsson et al., 2010; Shao et al., 2016). Cornified envelope genes (*Lce* and *Sprr* family members) and serine protease genes (*Serpinb3a* and *Serpinb3b*) were also deregulated in these animals, indicating epidermal barrier dysfunction (Fujimoto et al., 1997; Roberson et al., 2012; Shen et al., 2015). Meanwhile, increased *Krt6a*, *Krt16* and *Krt14* expression together with decreased *Krt15* and *Flg2* expression implicated abnormal keratinocyte differentiation in K5. Pp6^fl/fl^ mice (Alam et al., 2011; Lessard et al., 2013; Shen et al., 2015; Waseem et al., 1999). Conversely, elevated chemokine (*Cxcl1*, *Cxcl2* and *Cxcl5*) and cytokine (*Il1b*, *Il1f6*, *Il6*, *Il17a* and *Csf3*) production suggested the involvement of inflammatory reactions (**Figure 1I**).

The GSEA also indicated that the elevated and downregulated transcriptional signatures of K5. Pp6^fl/fl^ skin lesions significantly overlapped with DEGs identified in 92 human psoriatic skin samples relative to 82 normal skin samples (**Figure 1J**). The overlaps between the mouse transcriptional profile and the gene sets from human patients with atopic dermatitis were much weaker than the overlaps elicited by psoriatic DEGs (**Figure S2O**). Moreover, pathway enrichment analysis highlighted that TNF and IL-17 signaling is activated in the lesional skin of K5. Pp6^fl/fl^ mice (**Figure 1K**). These lesions responded well to cortisol (Halometasone) and anti-IL-17A treatment, demonstrating the active participation of the immune system, particularly IL-17A-producing cells, in the disease (**Figures S2P** and **1L**). Semi-quantitative cytokine array further confirmed the elevation of IL-23-T17-associated cytokines in K5. Pp6^fl/fl^ skin, as compared with Pp6^fl/fl^ controls, including IL-23, IL-12p40, IL-22, IL-20, IL-6, TNF-α, IL-17A and IL-17F (**Figure 1M**). Finally, scRNAseq on skin samples derived from diseased K5. Pp6^fl/fl^ mice found that Il17a and Il17f were expressed by both Trac^+^ αβT and Trdc^+^ γδT cells, but mainly by γδT cells in the diseased K5. Pp6^fl/fl^ skin (**Figure 1N**). Collectively, these histological, clinical and gene expression changes observed in K5. Pp6^fl/fl^ skin lesions are consistent with the hallmarks of psoriasis. Impaired PP6 expression in keratinocytes thus seems to be a pivotal event in psoriasis development.

### PP6 ablation promotes *ARG1* transcription through C/EBP-β activation in psoriatic keratinocytes

We next studied the molecular basis of the dysregulation in keratinocytes as a result of PP6 ablation (**Figure 2A**). Of note, genes deregulated in the absence of Pp6 showed a statistically significant overlap with CCAAT/enhancer-binding protein (C/EBP) target genes (**Figure 2B**). Specifically, various known C/EBPβ target genes, including *S100a8*/9, *Arg1*, *Il1f6*, *Tnf*, *Ccl20* and *Egr1,* were markedly upregulated (**Figure 2C**). In contrast to Pp6^fl/fl^ mice, we observed enhanced phospho-C/EBPβ (p-C/EBPβ, Thr-188) in the normal (unaffected) skin derived from K5. Pp6^fl/fl^ mice, and even higher levels in the lesional skin (**Figure S3A**). We noted the same phenomenon in human skin (**Figure S3B**). Total C/EBPβ was increased in the absence of Pp6 in the epidermis, in both non-lesional and lesional skin (**Figure S3C**). Importantly, the human psoriatic skin sections showed the identical C/EBP-β expression pattern to that observed in skin sections from diseased K5. Pp6^fl/fl^ mice (**Figure S3D**), with psoriatic skin expressing increased levels of C/EBP-β target genes as suggested by GSEA (**Figure 2D**). We confirmed C/EBP-β signaling activation in primary mouse keratinocytes from PP6-deficient mice, and in normal human epidermal keratinocytes (NHEKs) in which we knocked down PP6 by siRNA (**Figures S3E** and **S3F**). We also found that C/EBP-β co-immunoprecipitated with PP6 in the nuclear lysates of HaCaT keratinocytes (**Figure S3G**), suggesting that PP6 directly or indirectly interacted with C/EBP-β in nuclei.

**Figure 2.**
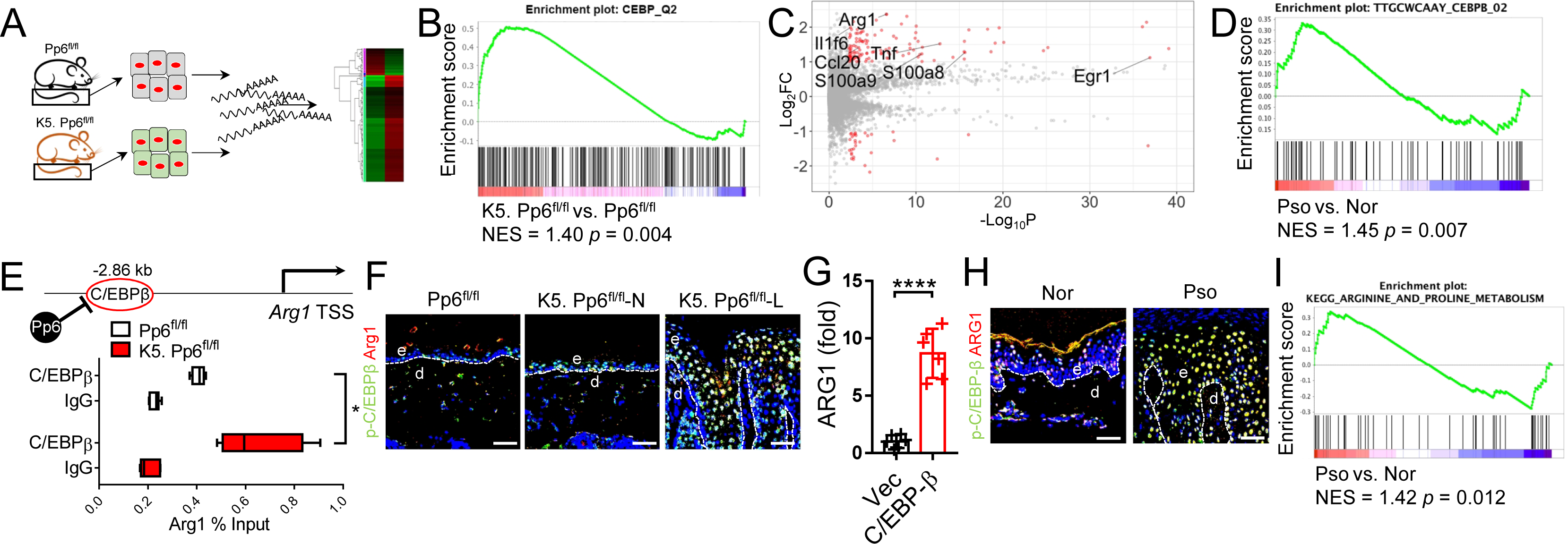
PP6 deficiency promotes C/EBP-β-mediated *ARG1* transcription in psoriatic keratinocytes. (**A**) Cultured murine keratinocytes derived from Pp6^fl/fl^ and K5. Pp6^fl/fl^ mouse tails were subjected to RNA-seq. (**B**) Transcriptional comparison of K5. Pp6^fl/fl^ and Pp6^fl/fl^ keratinocytes and enrichment of CEBP downstream signatures. (**C**) Volcano plot of the K5. Pp6^fl/fl^ versus the Pp6^fl/fl^ keratinocyte gene expression profiles. C/EBPβ target genes are indicated. (**D**) GSEA of DEGs in human psoriatic skin (GSE54456) with C/EBP-β target signatures enriched. (**E**) qPCR detection of the abundance of the *Arg1* promoter that immunoprecipitated with C/EBPβ in primary murine keratinocytes derived from Pp6^fl/fl^ and K5. Pp6^fl/fl^ mice (n = 5). The signals are presented as a percent of the total input chromatin. The boxes show the 25^th^ to 75th percentiles; the whiskers show the minimum to maximum; and the lines show the medians. (**F**) Immunofluorescent labelling of p-C/EBPβ and Arg1 in mouse skin samples. (**G**) qPCR detection of ARG1 in HaCaT cells transfected with an empty vector or a C/EBP-β overexpression plasmid (n = 6). The data are presented as the ratio of ARG1 to GAPDH relative to that in vector-transfected HaCaT cells. (**H**) Immunofluorescent labelling of p-C/EBP-β and ARG1 in human skin samples. (**I**) GSEA of DEGs in human psoriatic skin (GSE54456) with Arginine and Proline metabolism-associated genes enriched. For (**F** and **H**), the dotted lines indicate the border between the epidermis and the dermis; e, epidermis; d, dermis; Pp6^fl/fl^, healthy skin sample from control mice; K5. Pp6^fl/fl^-N, unaffected skin from K5. Pp6^fl/fl^ mice; K5. Pp6^fl/fl^-L, affected skin from K5. Pp6^fl/fl^ mice; Nor, normal human skin; Pso, psoriatic human skin; scale bars represent 50 μm. Data (**F** and **H**) are representative of > 5 samples in each group. Data (**E** and **G**) are representative of three independent experiments. \**p* < 0.05, \*\*\*\**p* < 0.0001, two-tailed Student’s *t*-test (means ± SEM).

We next generated PP6^-/-^ 293FT cells and investigated the C/EBP-β phosphorylation status in the absence or presence of PP6. Thr-179, Ser-184 and Thr-188 were hyper-phosphorylated in PP6^-/-^ 293FT cells (**Figure S3H**). PP6 shows de-phosphorylation specificity on “Thr-Pro” and “Ser-Pro” motifs (Rusin et al., 2015), and both Ser-184 and Thr-188 of C/EBP-β meet such criteria. We then generated plasmids expressing WT C/EBP-β, C/EBP-β^S184A^ and C/EBP-β^T188A^ and used Proximity Ligation Assay (PLA) to investigate the interactions between PP6 and the various C/EBP-β proteins. We found that PP6 interacted with WT C/EBP-β in cell nuclei (**Figure S3I**). Both PP6-C/EBP-β^S184A^ and PP6-C/EBP-β^T188A^ showed weakened interactions, and C/EBP-β with double mutations of Ser-184 and Thr-188 showed a cumulatively weak interaction with PP6 (**Figure S3I**). These data suggest that both C/EBP-β Ser-184 and Thr-188 are sites that are directly dephosphorylated by PP6 and the activation of C/EBP-β in both K5. Pp6^fl/fl^ mouse keratinocytes and human psoriatic keratinocytes could be a direct consequence of PP6 deficiency. Given that C/EBP-β is a key transcriptional factor downstream of the IL-17 receptor-mediated signaling (Song and Qian, 2013), we further investigated whether IL-17 caused PP6 downregulation in keratinocytes. PP6 was markedly reduced in NHEKs treated with IL-17A compared to untreated NHEKs (**Figure S3J**).

One of the most abundantly elevated genes in Pp6-deficient keratinocytes was *Arg1,* which encodes a key enzyme involved in the urea cycle that catalyzes Arginine hydrolysis to Ornithine and Urea. Human *ARG1* is highly expressed in the skin and a previous study found that *ARG1* ranked first in psoriasis-specific DEGs when compared with DEGs in other skin-confined diseases (Fagerberg et al., 2014; Swindell et al., 2016). Despite these findings, the functional significance of *ARG1* in psoriasis is unclear. Here, we found C/EBPβ enrichment in the *Arg1* promoter region in K5. Pp6^fl/fl^ keratinocytes as compared with Pp6^fl/fl^ keratinocytes (**Figure 2E**). Arg1 co-localized with p-C/EBPβ in the nuclei of suprabasal keratinocytes in the mouse epidermis, and elevated Arg1 expression correlated with C/EBPβ activation in both the Pp6-deficient non-lesional and lesional epidermis of K5. Pp6^fl/fl^ mice; this correlation was not evident in Pp6^fl/fl^ controls (**Figure 2F**). C/EBP-β overexpression in HaCaT keratinocytes led to a 8.49-fold increase in *ARG1* transcription (**Figure 2G**). Consistent with previous reports (Abeyakirthi et al., 2010; Bruch-Gerharz et al., 2003; Schnorr et al., 2005), we also confirmed ARG1 upregulation in the psoriatic epidermis compared to healthy donors (**Figure 2H**). More importantly, the DEGs that we identified as increased in the psoriatic epidermis overlapped with genes associated with Arginine and Proline metabolism (**Figure 2I**). Together, these data demonstrate that a PP6 deficiency in psoriatic keratinocytes facilitates C/EBP-β activation via phosphorylation of Ser-184 and Thr-188 sites of C/EBP-β and leads to increased ARG1 expression, which might further affect Arginine metabolism and urea cycle in psoriatic skin.

### Basal keratinocytes in the psoriatic epidermis exhibit *de novo* Arginine synthesis

We next aimed to detect the abundance of the substrates and products of ARG1 in human psoriatic samples using a QqQ-based LC-SRM-MS/MS metabolite quantitation system. Consistent with a previous metabolomics study (Kang et al., 2017), we detected an increased abundance of Urea in the sera of patients with psoriasis compared to that in healthy controls (**Figure 3A**). The serum Arginine and Ornithine levels were comparable between the two groups (**Figure 3A**). Because we showed that the ARG1 protein levels were increased in the psoriatic epidermis (**Figure 2H**), we hypothesized that arginase activity is enhanced in the psoriatic epidermis while a compensatory mechanism likely exists to support local Arginine consumption. To confirm our hypothesis, we generated scRNAseq datasets from samples of the psoriatic epidermis and partitioned the cells according to their countermarks (**Figure 3B**). We found that basal keratinocytes (labeled by KRT14, KRT15 and CXCL14) uniquely expressed the Arginine biosynthesis rate-limiting enzyme, Argininosuccinate synthetase 1 (ASS1) (**Figure 3C**). By contrast, these cells did not show specific expression of the Arginine transporters SLC7A1, SLC7A2, SLC7A5, SLC7A7 or SLC3A2 (**Figure 3C**). Compared to the healthy epidermis, the psoriatic epidermis showed augmented ASS1-expressing region (**Figure 3D**). We then generated merged scRNAseq datasets of three psoriatic epidermis samples and three healthy epidermis samples to determine “KRT14”, “KRT5”, “KRT10” and “ASS1” expression patterns. We found that ASS1 was strictly expressed in KRT14^hi^KRT5^hi^KRT10^lo^ basal keratinocytes (**Figure 3E**). We then calculated the “ASS1 score” [(ASS1 average expression value of ASS1^+^ keratinocytes) × (ASS1^+^ keratinocyte number) / (total keratinocyte number)] for each sample to determine the contribution of *de novo* Arginine synthesis to Arginine compensation in the epidermis. The psoriatic keratinocytes had a higher ASS1 score than the control keratinocytes (**Figure 3F**), which is suggestive of a faster urea cycle that refreshes the psoriatic epidermis.

**Figure 3.**
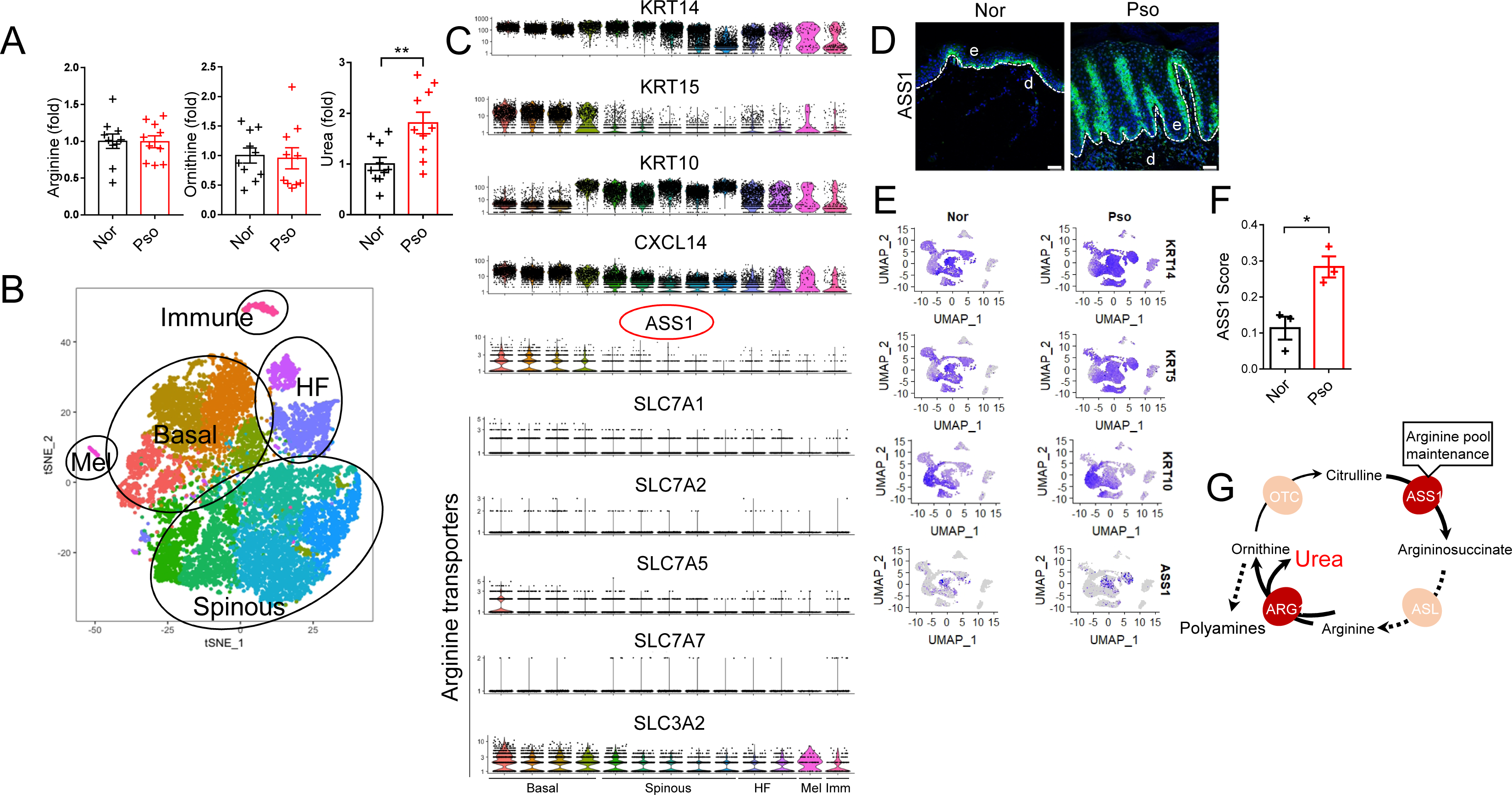
De *novo* Arginine synthesis is featured in basal keratinocytes of psoriatic epidermis. (**A**) Relative quantitation of Arginine, Ornithine and Urea levels in the sera of patients with psoriasis and healthy donors (n = 10). (**B**) Representative t-SNE plot of scRNAseq of epidermal cells derived from a patient with psoriasis (n = 3). Basal keratinocyte, spinous keratinocyte, hair follicle (HF), melanocyte (Mel) and immune cell (immune) compartments are encircled. (**C**) Violin plots of the indicated gene expression levels distributed across distinct cell compartments. (**D**) Immunofluorescent labelling of ASS1 in skin sections derived from healthy donors and patients with psoriasis. The data are representative of > 5 samples in each group. The dotted lines indicate the border between the epidermis and the dermis; e, epidermis; d, dermis; Nor, normal human skin; Pso, psoriatic human skin; scale bars represent 50 μm. (**E**) Feature-plots of indicated genes in merged scRNAseq datasets of three healthy epidermal samples and three psoriatic epidermal samples. (**F**) ASS1 score of keratinocytes from 3 healthy donors and 3 patients with psoriasis, calculated using the formula: (ASS1 average expression value of ASS1^+^ keratinocytes) × (ASS1^+^ keratinocyte number) / (total keratinocyte number). (**G**) Urea cycle reprogramming in psoriatic keratinocytes. \**p* < 0.05, \*\**p* < 0.01, two-tailed Student’s *t*-test (means ± SEM).

We also used scRNAseq to study Arginine compensation mechanisms in Pp6-deficient murine keratinocytes. We partitioned the keratinocytes according to a previous single-cell transcriptome study of the mouse epidermis, where interfollicular (IFE) basal keratinocytes were marked as Krt14^hi^Mt2^hi^ (Joost et al., 2016). IFE basal keratinocytes from both control and K5. Pp6^fl/fl^ mice highly and exclusively expressed Ass1, while the Arginine transporter genes were either undetected or showed no cell specificity (**Figure S4**). We thus conclude that *de novo* Arginine synthesis is also a feature of mouse IFE basal keratinocytes. The proportion of Ass1^+^ cells was increased in the Pp6-deficient epidermis, which suggests that more keratinocytes in this context have Arginine-producing capacity than keratinocytes in the WT epidermis. This effect is likely to support quick Arginine consumption. Taken together, psoriatic keratinocytes rely on *de novo* Arginine biosynthesis to sustain increased ARG1 activity and the subsequent generation of downstream metabolites (**Figure 3G**).

### Urea cycle rewiring is characteristic of Pp6-deficient keratinocytes

Arginase and nitric oxide synthase (NOS) share Arginine as a common substrate. We found that Arg1 augmentation in K5. Pp6^fl/fl^ keratinocytes results in a 50% reduction in cellular NO production compared with Pp6^fl/fl^ controls (**Figure 4A**). Analogously, we found that psoriatic skin produces an impaired transcriptional response to NO compared to normal skin (**Figure 4B**). To better understand the metabolic profile of K5. Pp6^fl/fl^ keratinocytes *in vivo*, we performed an unbiased UPLC-QTOF/MS-based metabolomics analysis of the epidermis derived from K5. Pp6^fl/fl^ and Pp6^fl/fl^ mice. The two groups showed unique metabolic profiles as indicated by the principal component analysis (PCA; **Figure 4C**). Specifically, direct comparison of the metabolite levels between the two groups revealed an enrichment in fatty acids including Stearic, Oleic and Palmitic acid derivatives in the epidermis of K5. Pp6^fl/fl^ mice (**Figure 4D**), and an increase in TCA cycle-anaplerotic reacting amino acids including L-Glutamine, L-Glutamic acid and L-Aspartic acid compared to Pp6^fl/fl^ controls (**Figure 4E**). In parallel with this molecular rewiring, we found that L-Ornithine, the direct Arg1 product, and Ornithine downstream products (Citrulline, Proline, Creatine and Spermidine) were elevated in the epidermis of K5. Pp6^fl/fl^ mice compared to littermate controls (**Figure 4E**). QqQ-based LC-SRM-MS/MS also revealed that the urea levels were 1.88-fold higher in the K5. Pp6^fl/fl^ epidermis and 1.2-fold higher in the K5. Pp6^fl/fl^ sera compared to Pp6^fl/fl^ mice (**Figures 4F** and **4G**). Combining the transcriptome and metabolomics data, we found that urea cycle reprogramming was a hallmark of K5. Pp6^fl/fl^ keratinocytes (**Figures 4H** and **4I**).

**Figure 4.**
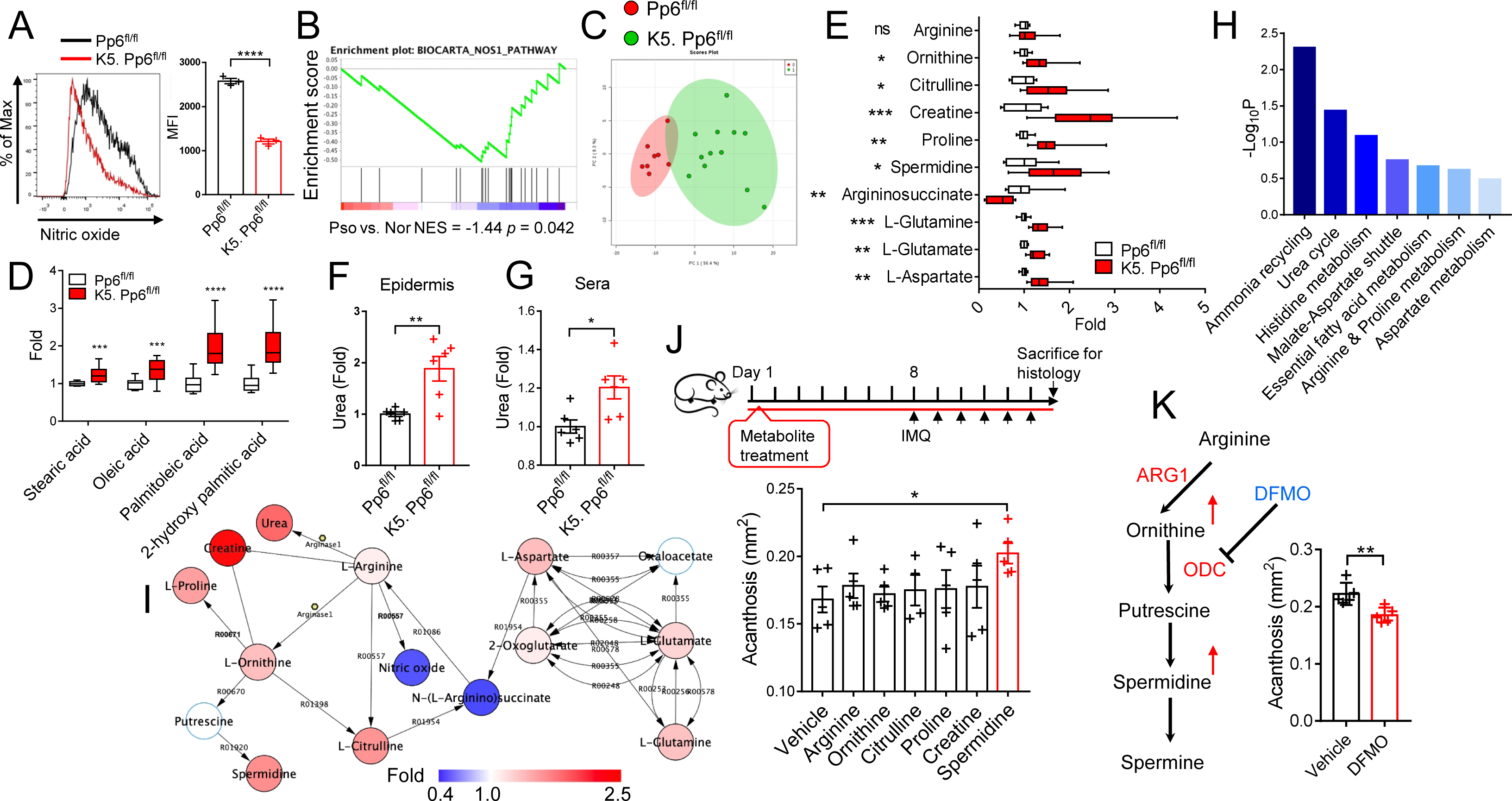
The urea cycle undergoes rewiring in Pp6-deficient keratinocytes. (**A**) Flow cytometric analysis of NO levels in Pp6^fl/fl^ and K5. Pp6^fl/fl^ keratinocytes (n = 3). (**B**) GSEA of DEGs in human psoriatic skin (GSE54456) with NO-responsive genes enriched. (**C**) PCA of the epidermal metabolites derived from Pp6^fl/fl^ (n = 8) and K5. Pp6^fl/fl^ (n = 12) mice. (**D**) Relative quantification of the indicated fatty acids extracted from the epidermis of Pp6^fl/fl^ and K5. Pp6^fl/fl^ mice. (**E**) Relative quantitation of urea cycle-associated metabolites based on a metabolomics analysis of the K5. Pp6^fl/fl^ and Pp6^fl/fl^ epidermis. The boxes show the 25^th^ to 75th percentiles; the whiskers show the minimum to maximum; and the lines the show medians. (**F** and **G**) Relative quantitation of Urea levels in the epidermis (**F**) and sera (**G**) of Pp6^fl/fl^ and K5. Pp6^fl/fl^ mice (n = 6). (**H**) Quantitative enrichment analysis of the metabolites derived from Pp6^fl/fl^ and K5. Pp6^fl/fl^ mice using MetaboAnalyst 3.0. (**I**) Urea cycle-associated metabolites altered with Pp6 ablation in the epidermis of K5. Pp6^fl/fl^ mice, relative to those of Pp6^fl/fl^ controls. The fold changes of the metabolites are indicated by the continuous color. Hollow circles with light blue edges indicate the undetected metabolites. KEGG reaction numbers are marked between the contiguous metabolites. (**J**) Degree of acanthosis of IMQ-induced psoriasis-like skin inflammation treated with the indicated urea cycle-associated metabolites (n = 4∼5) according to the indicated study protocol (top panel). (**K**) Acanthosis of IMQ-induced psoriasis-like skin inflammation treated with vehicle or DFMO (n = 5); the function of DFMO is illustrated in the left-hand panel. The data (**A**, **J** and **K**) are representative of three independent experiments. \**p* < 0.05, \*\**p* < 0.01, \*\*\**p* < 0.001, \*\*\*\**p* < 0.0001, two-tailed Student’s *t*-test (means ± SEM).

We then treated IMQ-induced mice with urea cycle-associated metabolites to study their respective effects on psoriasis-like skin inflammation. Among all the tested metabolites, only Spermidine significantly exacerbated epidermal hyperplasia (**Figure 4J**). Correspondingly, the Ornithine decarboxylase (ODC) inhibitor DL-2-(Difluoromethyl)-ornithine hydrochloride (DFMO), which restricts polyamine generation, mitigated IMQ-induced skin inflammation (**Figure 4K**). Taken together, we propose that loss of Pp6 promotes transcriptional and coordinated metabolic adaptations in keratinocytes centered on urea cycle rewiring. Spermidine — a polyamine branched from the urea cycle — is a pro-inflammatory factor in psoriasis-like skin inflammation.

### Pp6-deficient keratinocytes show enhanced oxidative phosphorylation

NO negatively impacts OXPHOS efficiency by interacting with the mitochondrial Complex I and cytochrome c oxidase (Complex IV) (Sarti et al., 2012)(**Figure S5A**). We next decided to investigate the OXPHOS status of Pp6^fl/fl^ and K5. Pp6^fl/fl^ keratinocytes, as the latter showed defective NO production (**Figure 4A**). To do so, we measured the keratinocyte oxygen consumption rate (OCR) after treatment with different electron transport chain (ETC) modulators. As expected, Pp6 deficiency in keratinocytes led to an elevation in the basal respiration level and spare respiratory capacity despite a decrease in non-mitochondrial respiration **(Figure S5B**). Meanwhile, we identified extended mitochondrial fuel flexibility for glutamine and glucose in K5. Pp6^fl/fl^ keratinocytes, as indicated by the different dependencies on, and capacities to consume, both fuels **(Figure S5C**). Conversely, K5. Pp6^fl/fl^ keratinocytes showed a subtle decrease in their glycolytic capacity (**Figure S5D**). We also found that ATP production by K5. Pp6^fl/fl^ keratinocytes was increased 1.24-fold compared to Pp6^fl/fl^ controls, which was attributed to enhanced OXPHOS (**Figure S5E**).

We then detected the protein levels of ETC components in murine and human samples. In mice, we found that Complex II (Succinate dehydrogenase complex, subunit A, Sdha) and Complex IV (cytochrome c oxidase II, mt-Co2 and cytochrome c oxidase subunit 4I1, Cox4i1) were both enriched in K5. Pp6^fl/fl^ keratinocytes compared to Pp6^fl/fl^ controls, supporting the functional phenotype of Pp6-deficient keratinocytes (**Figure S5F**). In human samples, we only detected COX4I1 in the basal layer of the epidermis of healthy skin, while the psoriatic epidermis showed condensed and extended COX4I1 expression (**Figure S5G**).

We finally treated IMQ-induced mice with the OXPHOS inhibitor Oligomycin. Here, we found that topical Oligomycin application significantly alleviated the psoriasis-like phenotype in C57BL/6 mice, as indicated by delayed ear swelling and relieved acanthosis (**Figures S5H-5J)**. These data suggest that OXPHOS, as a deregulated process in keratinocytes with Pp6 deficiency, might be a target for psoriasis therapy.

### Excessive polyamine production by psoriatic keratinocytes enables self-RNA sensing by myeloid DCs

Thus far, we have shown that arginase is increased in Pp6-deficient keratinocytes, which endows psoriatic keratinocytes with urea cycle rewiring and enhanced OXPHOS; however, key questions remained as to whether and how the identified metabolic adaptions within psoriatic keratinocytes affects the immune response.

Previous studies unveiled dramatic increases in the levels of polyamines, including Putrescine, Spermine and Spermidine, in the skin and blood of patients with psoriasis compared to those of healthy donors (Broshtilova et al., 2013; Proctor et al., 1975). Because we showed in this study that polyamine supplementation is detrimental in a mouse model of psoriasis (**Figure 4J**), we next aimed to characterize the possible underlying mechanisms of their effect on skin inflammation. Polyamines are highly charged aliphatic polycations that bind and aggregate nucleic acids, rendering their application in gene transfection (Zavradashvili et al., 2019), which resembles self-RNA internalization in principle. Given that self-RNA internalization is a pivotal pathogenic factor of psoriasis and other chronic inflammatory disorders (Celhar et al., 2015; Ganguly et al., 2009), we hypothesized that polyamines might facilitate this process in the context of disease. To test this, we stimulated secondary lymphoid organ-derived DCs with supernatants of UV-irradiated murine keratinocytes, which underwent apoptosis and necrosis with the release of self-nucleotides (Lovgren et al., 2004). Supernatants from K5. Pp6^fl/fl^ keratinocytes showed a higher potency to activate DCs than supernatants from Pp6^fl/fl^ keratinocytes; this difference was abrogated when K5. Pp6^fl/fl^ keratinocytes were pre-treated with DFMO for 48 h to eliminate intracellular polyamines or when the dead keratinocyte supernatants were pre-treated with RNase (**Figure 5A**). These results indicate that polyamines are possibly involved in sensing of keratinocyte-derived self-RNA by DCs.

**Figure 5.**
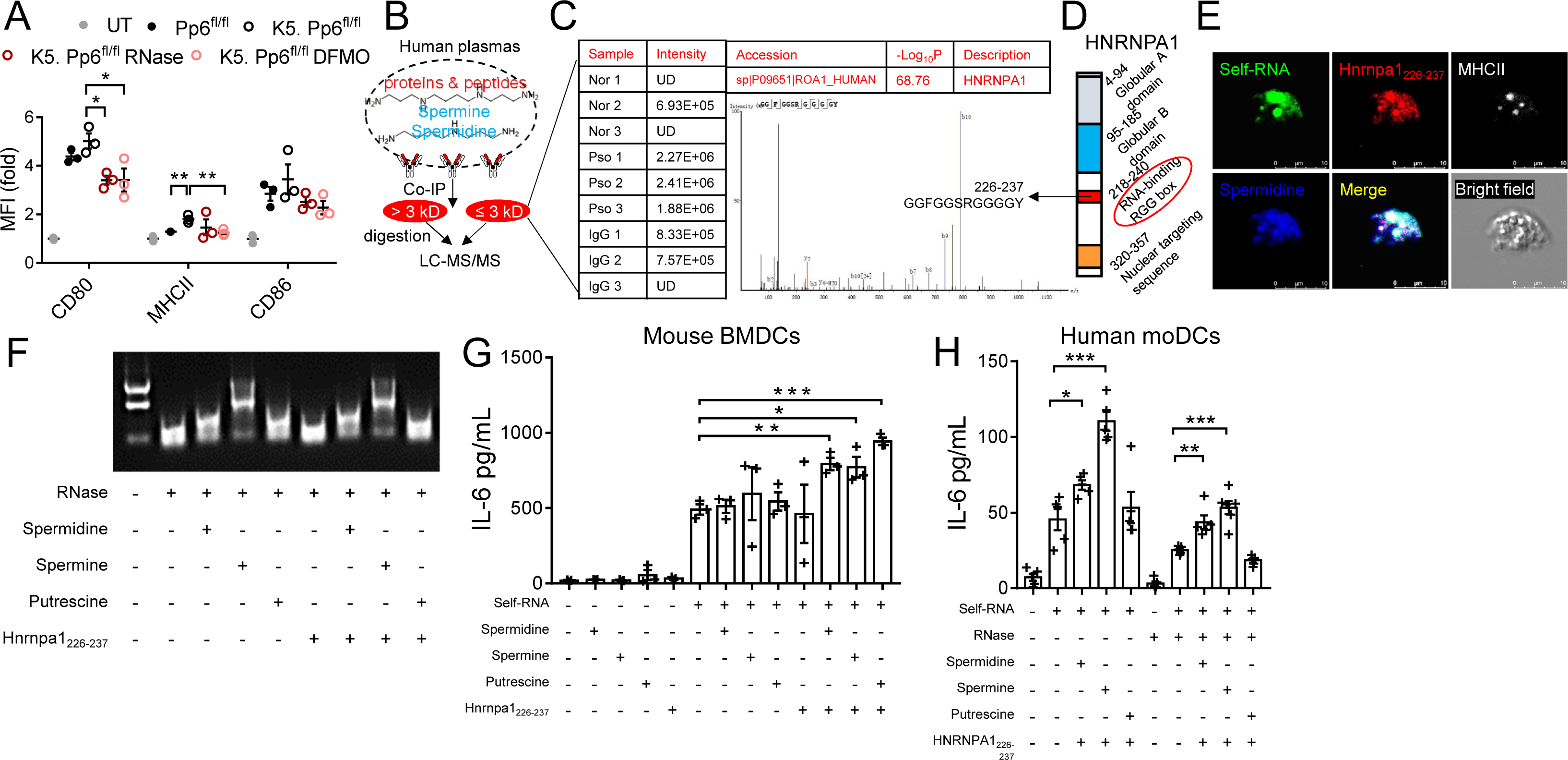
Polyamines enable self-RNA sensing by myeloid DCs. (**A**) Statistics of the flow cytometric analysis of DCs cultured overnight in dead keratinocyte supernatants generated from Pp6^fl/fl^ and K5. Pp6^fl/fl^ mice (n = 3). (**B**) Study protocol to identify proteins or peptides in a complex with Spermidine or Spermine in human blood plasma samples. (**C**) LC-MS/MS-based detection of the abundance of HNRNPA1226-237 in the immunoprecipitates pulled by anti-Spermidine/Spermine from the blood plasma of patients with psoriasis (n = 3, Pso) and healthy donors (n = 3, Nor). Immunoprecipitates pulled by rabbit IgG from the blood plasma of patients with psoriasis (n = 3, IgG) served as negative controls. The MS chromatogram verifies the presence and sequence of HNRNPA1226-237. UD, undetected. (**D**) The human HNRNPA1 protein functional domains described in UniProt. (**E**) Confocal microscopic images of mouse BMDCs treated with 488-labeled RNA, TMR-labeled Hnrnpa1226-237 and 2-(methylamino) benzoic acid-labeled Spermidine, with the cells co-stained with MHCII. (**F**) Agarose gel showing the degradation of RNA treated with the indicated reagents. (**G**) ELISA quantification of IL-6 levels in mouse BMDC cultures stimulated with the indicated reagents (n = 3). (**H**) ELISA quantification of IL-6 in human moDC cultures stimulated with the indicated reagents (n = 5). The data (**A** and **E**-**H**) are representative of three independent experiments. \**p* < 0.05, \*\**p* < 0.01, \*\*\**p* < 0.001, two-tailed Student’s *t*-test (means ± SEM).

However, we found that polyamines alone could not achieve the intracellular delivery of RNA molecules to cells (**Figure S6**). We assumed that cargo molecules in complex with polyamines and RNAs might assist with self-RNA internalization in psoriasis. Using an antibody against Spermidine and Spermine, we immunoprecipitated peptides ≤ 3 kilodalton (kD) and proteins > 3 kD that were in a complex with Spermidine or Spermine in the plasmas of patients with psoriasis and healthy donors (**Figure 5B**). We identified a 12-amino acid peptide (GGFGGSRGGGGY) originating from heterogeneous nuclear ribonucleoprotein A1 (HNRNPA1) that was enriched in the immunoprecipitates from the patients (**Figure 5C**). This peptide (designated as HNRNPA1226-237) contains an RGG-box that is indispensable for RNA binding of HNRNPA1, which is a probable psoriasis autoantigen (Jones et al., 2004) (**Figure 5D**). Using an established Spermidine labeling strategy (Wolf et al., 2006), we were able to observe the co-localization of exotic Spermidine with 488-labeled self-RNA and Tetramethylrhodamine (TMR)-labeled Hnrnpa1226-237 within mouse bone marrow-derived DCs (BMDCs). This unmitigated co-localization of these molecules together with murine MHCII informed us about not only the interaction between self-RNA-polyamine-Hnrnpa1226-237 complexes, but also about the immunogenicity of these complexes (**Figure 5E**). These self-RNA–polyamine–Hnrnpa1226-237 complexes were more internalized by BMDCs than self-RNA alone (**Figure S6**), suggestive of increased intracellular delivery of self-RNA to BMDCs.

By looking into the RNA-seq data of human primary keratinocytes, we also found that the RNA degradation pathway is highly activated in keratinocytes derived from both the uninvolved and involved skin of patients with psoriasis compared with keratinocytes from healthy skin (**Figure S7A**). Meanwhile, the widespread existence of extracellular RNases restricts the internalization of self-RNA from dying keratinocytes by DCs. Although polyamines supposedly stabilize RNA, their ability to protect self-RNA from nuclease degradation has not been previously investigated. Here, we found that polyamines protect self-RNA from RNase digestion to varying degrees independent of the presence or absence of Hnrnpa1226-237 (**Figure 5F**). Notably, Spermine showed superior protective capability over Spermidine and Putrescine, while Hnrnpa1226-237 alone could not prevent RNA degradation when digested with RNase (**Figure 5F**).

We next detected cytokine production by BMDCs in different settings of stimulation to investigate whether polyamines indeed enabled self-RNA sensing by DCs. Exposure to polyamines or Hnrnpa1226-237 alone could not activate BMDCs (**Figures 5G** and **S7B**) while polyamine–self-RNA or Hnrnpa1226-237–self-RNA complexes induced equivalent levels of IL-6 and Tnf production by BMDCs compared to self-RNA alone (**Figures 5G** and **S7B**). We observed significantly increased IL-6 and Tnf production when stimulating BMDCs with self-RNA–polyamine–Hnrnpa1226-237 complexes (**Figures 5G** and **S7B**). Similarly, we found that self-RNA–polyamine–Hnrnpa1226-237 complexes could stimulate human monocyte derived-DCs (moDCs) to produce IL-6, even when the complexes were treated with RNase prior to stimulation (**Figure 5H**). Interestingly, the different polyamines showed distinct stimulatory efficiencies between murine BMDCs and human moDCs, and moDCs did not produce considerable TNF-α upon RNA-containing complex stimulation except for self-RNA–Spermine–HNRNPA1226-237 (**Figure S7C**). Together with Hnrnpa1226-237, polyamines did not promote DNA sensing by plasmacytoid DCs (pDCs, **Figure S7D**). Although we did not identify clonal expansion when naïve CD4^+^ T cells were co-cultured with BMDCs pre-stimulated with self-RNA–polyamine–Hnrnpa1226-237 complexes (**Figure S7E**), the percentage of Th17 cells increased in a co-culture system of BMDCs pre-stimulated with self-RNA–Putrescine–Hnrnpa1226-237 complexes (**Figure S7F**).

We next investigated whether there existed other cargo peptides that could assist with self-RNA sensing processes. Importantly, we detected a Keratin 10-derived peptide, Krt1017-28 (SGGGGGGGSVRV) in the sera of K5. Pp6^fl/fl^ but not Pp6^fl/fl^ mice (**Figure S7G**). Similar to HNRNPA1226-237, Krt1017-28 is positively charged at pH 7 and like many RNA-binding proteins, is glycine rich (Mousavi and Hotta, 2005). We next used self-RNA–Krt1017-28–polyamine complexes to stimulate BMDCs and found that BMDCs secreted IL-6 upon stimulation (**Figure S7H**). Taken together, we conclude that accumulated polyamines from hyper-proliferative keratinocytes promote self-RNA sensing by DCs with the assistance of cargo peptides.

### Self-RNA sensing facilitated by polyamines depends on endosomal Tlr7 in myeloid DCs

TLR7 and TLR8 are pivotal for DCs to sense exogenously delivered RNA molecules in humans (Sioud, 2006), but TLR8 is non-functional in mice (Hemmi et al., 2002). We next tested whether self-RNA–polyamine–Hnrnpa1226-237 sensing by murine BMDCs is Tlr7 dependent by using Tlr7^-^ mice. We found that IL-6 production (as an indicator of DC activation) was agitated in WT BMDCs stimulated with self-RNA-polyamine-Hnrnpa1226-237 complexes, but restored in Tlr7-deficient BMDCs treated with self-RNA-polyamine-Hnrnpa1226-237 complexes (**Figure 6A**). Because polyamines protect RNA from degradation *in vitro* (**Figure 5F**), we questioned whether RNA molecules in complex with polyamines internalized by DCs are protected and retained in endosomes. We thus treated murine BMDCs with self-RNA–polyamine–Hnrnpa1226-237 complexes containing total RNA derived from human HaCaT cells for 4 h. We then extracted endosomal RNA from these BMDCs and measured the human 28S rRNA abundance. We found an elevated level of exogenous RNA molecules retained in the BMDC endosomes when the RNAs were in a complex with a polyamine and Hnrnpa1226-237 compared to when they were free (**Figure 6B**). IkappaB-ζ is an essential transcription factor involved in Tlr7-downstream signaling and is required for IL-6 generation (Yamamoto et al., 2004). We found that IkappaB-ζ was augmented in the nuclei of BMDCs stimulated with self-RNA–polyamine–Hnrnpa1226-237 complexes (**Figure 6C**).

**Figure 6.**
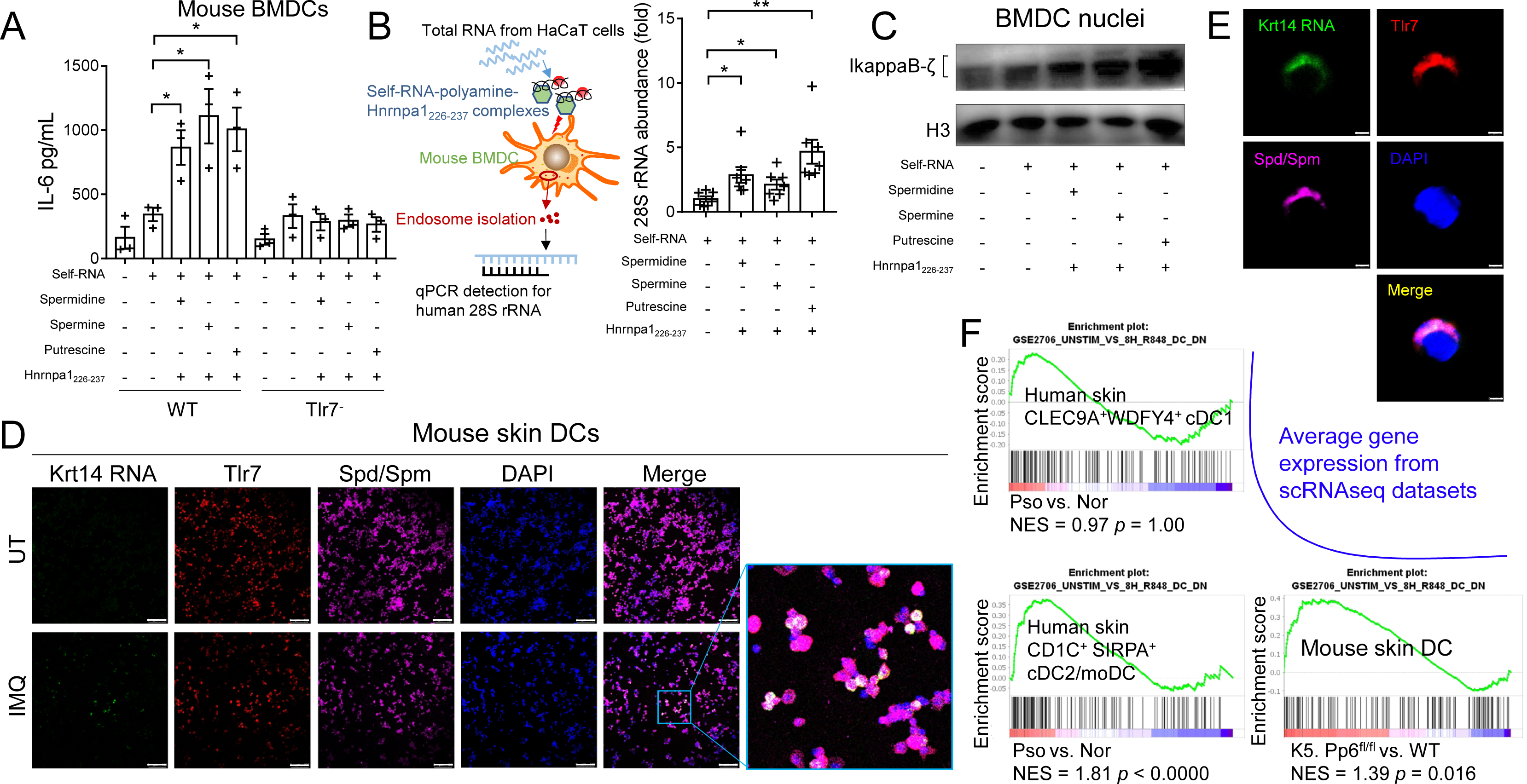
Polyamine-mediated self-RNA sensing requires endosomal Tlr7. (**A**) ELISA quantification of IL-6 in WT and Tlr7-deficient BMDC cultures stimulated with the indicated reagents (n = 3). (**B**) qPCR detection of human 28S rRNA in endosomes isolated from mouse BMDCs treated with the indicated reagents (n = 7). The fold changes relative to the self-RNA group are presented. (**C**) Immunoblot analysis of IkappaB-ζ expression in nuclear fractions of BMDCs treated with the indicated reagents for 4 h. (**D** and **E**) RNAscope-based FISH of Krt14 combined with immunofluorescent labelling of Tlr7 and Spermidine/Spermine on DCs isolated from mouse skin treated with IMQ or untreated (UT). Scale bars represent 100 μm (**D**) or 3 μm (**E**). (**F**) GSEA of all genes in CLEC9A^+^WDFY4^+^ cDC1 and in CD1C^+^ SIRPA^+^ cDC2/moDC from human psoriatic skin relative to healthy skin with the genes induced in R848 stimulated DCs enriched (GSE2706). GSEA of skin DCs in lesional K5. Pp6^fl/fl^ skin relative to normal mouse skin with the genes induced in R848 stimulated DCs enriched (GSE2706). The data (**A**-**E**) are representative of two independent experiments. \**p* < 0.05, \*\**p* < 0.01, two-tailed Student’s *t*-test (means ± SEM).

We next directly visualized the internalization of keratinocyte-derived RNA by DCs in psoriasis-like skin lesions. We performed RNAscope-based fluorescence *in situ* hybridization (FISH) to detect single molecule Krt14 mRNA (exclusively expressed by keratinocytes) combined with immunofluorescent labelling of Tlr7 and Spermidine/Spermine on DCs isolated from murine skin treated or not with IMQ. We detected clear Krt14 mRNA signals in DCs from IMQ-treated mouse skin, which co-localized with Tlr7 and Spermidine/Spermine (**Figures 6D** and **6E**). To gain a deeper insight into the *in vivo* DC status in psoriasis-like skin lesions, we merged and visualized scRNAseq datasets of DCs from skin samples of diseased K5. Pp6^fl/fl^ mice, untreated C57BL/6 mice (control) and IMQ-treated C57BL/6 mice. We found that DC populations in lesional K5. Pp6^fl/fl^ skin and in IMQ-treated skin were radically different from the DC populations in control skin (**Figures S8A** and **S8B**). Specifically, Lyz1 was highly expressed by DCs in untreated mouse skin; Cd301b (Mgl2)^+^Sirpa^+^Cd11b (Itgam)^+^ cDC2 cells were dominant in IMQ-treated skin; and Cd301b (Mgl2)^-^Sirpa^+^Cd11b (Itgam)^+^Mafb^+^ DCs (cDC2 or monocyte-derived DCs) were dominant in lesional K5. Pp6^fl/fl^ skin (**Figure S8C**). Further analysis of the scRNAseq datasets generated from dermal immune cells of patients with psoriasis and healthy donors showed that TLR7 signaling was activated in psoriatic CD1C^+^SIRPA^+^ cDC2 or moDC subsets, but not in the CLEC9A^+^WDFY4^+^ cDC1 subset (**Figure 6F**). Of note, Tlr7 signaling was activated in skin DCs from lesional K5. Pp6^fl/fl^ skin compared to normal dermal DCs (**Figure 6F**). Altogether, these data support that self-RNA–polyamine–Hnrnpa1226-237 complexes activate endosomal Tlr7 in myeloid DCs of psoriasiform skin lesions.

### Restoring the urea cycle in keratinocyte alleviates psoriasis-like skin inflammation

Based on the mechanistic information we had gathered thus far, we finally hypothesized that targeting key nodes of the reprogrammed keratinocyte metabolic network might improve psoriasis-like skin inflammation. Because increased Arg1 is the feature of psoriatic epidermis and the direct cause of urea cycle rewiring, we decided to use nor-NOHA to inhibit Arg1 enzymatic activity. We topically injected C57BL/6 mice with nor-NOHA (100 μg/mouse/day) and smeared 5% IMQ cream on the ear daily for 6 consecutive days. Inhibiting Arg1 delayed ear swelling and alleviated the inflammatory phenotype (erythema, thickness and scaling) in IMQ-treated mice compared to vehicle (PBS)-treated mice (**Figures 7A** and **7B**). Histological analysis showed that nor-NOHA remarkably suppressed epidermal hyperplasia (**Figures 7C** and **7D**). Mitigated DC activation as indicated by decreased expression levels of CD80, CD86 and MHCII on cell surface was detected in the dermis and draining lymph nodes of mice treated with nor-NOHA or DFMO compared to vehicle-treated mice on Day 3 of IMQ induction (**Figure 7E**). To better evaluate the translational potential of inhibiting arginase activity to treat psoriasis, we also exposed cynomolgus monkeys to IMQ on the back-shoulders to induce psoriasis-like skin inflammation; we then monitored the effects of nor-NOHA on inflammation. nor-NOHA treatment significantly reduced the clinical scores and acanthosis of the skin inflammation as well as the infiltration of CD4^+^ and CD11C^+^ cells, as compared with vehicle-treated controls (**Figures 7F**-**7J**).

**Figure 7.**
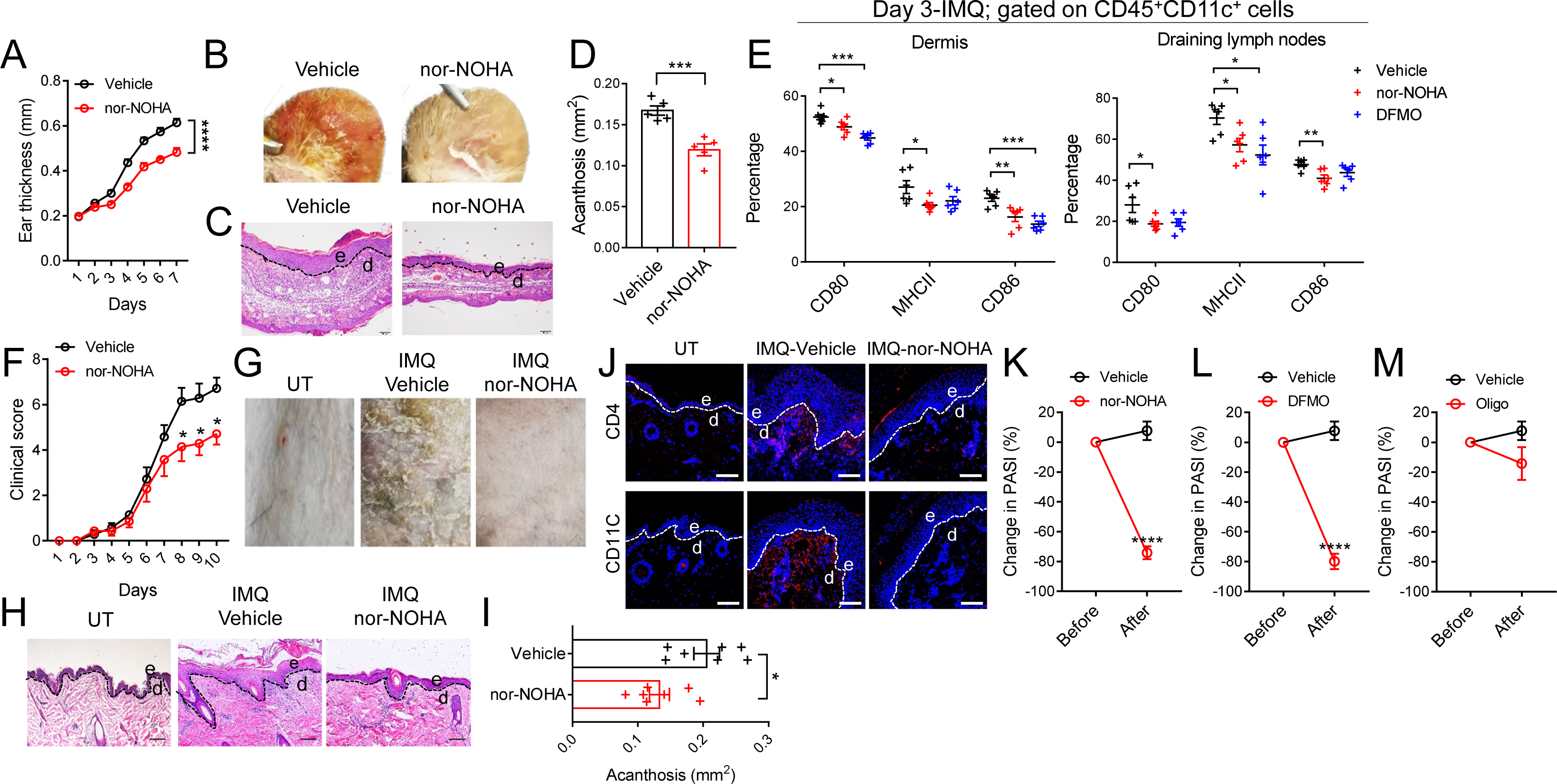
Targeting keratinocyte metabolic rewiring alleviates psoriasis-like skin inflammation in mice and non-human primates. (**A**) Phenotypic representation of mouse ears of IMQ-induced C57BL/6 mice treated with vehicle and nor-NOHA (n = 5). (**B**) Ear thickness changes of IMQ-induced C57BL/6 mice treated with vehicle and nor-NOHA. (**C**) Representative H&E staining of mouse ears of IMQ-induced C57BL/6 mice treated with vehicle and nor-NOHA. (**D**) Acanthosis of mouse ears of IMQ-induced C57BL/6 mice treated with vehicle and nor-NOHA. (**E**) Statistics of the flow cytometric analysis of dermal DCs and DCs in skin draining lymph nodes from 3-day-IMQ-induced C57BL/6 mice treated with vehicle, nor-NOHA or DFMO (n = 6). (**F**) Phenotypic representation of untreated (UT) and IMQ-induced Cynomolgus monkey skin treated with vehicle or nor-NOHA (n = 7). (**G**) Clinical score changes of IMQ-induced Cynomolgus monkey skin treated with vehicle or nor-NOHA. (**H**) Representative H&E staining of UT and IMQ-induced Cynomolgus monkey skin treated with vehicle or nor-NOHA. (**I**) Acanthosis of IMQ-induced Cynomolgus monkey skin treated with vehicle or nor-NOHA. (**J**) Representative CD4 and CD11C immunofluorescent labelling of UT and IMQ-induced Cynomolgus monkey skin treated with vehicle or nor-NOHA. Diseased K5. Pp6^fl/fl^ mice were treated with nor-NOHA (**K**), DFMO (**L**) or Oligomycin (Oligo, **M**) every other day for 2 weeks. Changes in PASI before and after treatment were calculated. For (**C**, **H** and **J)**, the dotted lines indicate the border between the epidermis and the dermis; e, epidermis; d, dermis; the scale bars represent 100 μm. The data (**A**-**E**) are representative of three independent experiments. The data (**F**-**J**) are representative of two independent experiments. The data (**K-M**) are representative of two independent experiments with two mice in each group. \**p* < 0.05, \*\*\**p* < 0.001, \*\*\*\**p* < 0.0001, two-way ANOVA (**A**) and two-tailed Student’s *t*-test (means ± SEM).

To test the effects of nor-NOHA, DFMO and Oligomycin, which inhibit arginase activity, polyamine synthesis and OXPHOS respectively, on the dispersed skin lesions of K5. Pp6^fl/fl^ mice, we injected these compounds intraperitoneally to the diseased mice every other day for 2 weeks. nor-NOHA and DFMO treatment significantly reduced the PASI of the diseased mice but Oligomycin had no notable therapeutic effects (**Figures 7K**-**M**). These data support that preventing urea cycle rewiring rescues formed skin lesions of K5. Pp6^fl/fl^ mice. OXPHOS might be a pathogenic factor of disease development but not maintenance in K5. Pp6^fl/fl^ mice.

## DISCUSSION

Pioneering genetic mouse models have used gene overexpression approaches to agitate the inflammatory response. By contrast, we used a knockout strategy in this study to break the intrinsic balance of epidermis homeostasis and achieve a predisposed disease status. Specifically, we deleted Pp6 in keratinocytes, which resulted in coordinated transcriptional and metabolic rewiring of urea cycle in keratinocytes. We also found that keratinocyte-derived polyamines trigger self-RNA endosomal sensing, and for the first time identified a direct link between keratinocyte hyperproliferation and innate immune responses in psoriasis maintenance. These findings provide a step forward in the mechanistic delineation of chronic inflammation. Targeting nodal points (arginase activity, polyamine generation and OXPHOS) in these metabolic adaptations elicited by Pp6 ablation might constitute a novel strategy to improve psoriasis and other chronic inflammatory diseases in affected patients (**Figure S8**).

In 2016, Swindell *et al*. performed a “cross-disease” RNA-seq meta-analysis on psoriasis and other skin-confined disorders (Swindell et al., 2016). Their findings led to the notion that psoriasis shares most DEGs with many other skin conditions while psoriasis-specific DEGs are expressed by keratinocytes and are induced by IL-17A (Swindell et al., 2016). Here, we show that IL-17A downregulates PP6 in keratinocytes and that Pp6-deficient keratinocytes harbor DEGs that overlap with genes that are transcriptionally activated by C/EBPβ, a key transcriptional factor involved in IL-17 receptor-mediated signaling (Song and Qian, 2013). Concomitantly, *ARG1* is highly expressed upon C/EBP-β activation due to PP6 ablation in keratinocytes; this gene was also the most psoriasis-specific gene identified in the previous “cross-disease” study. Together, these findings suggest that impaired PP6 expression in psoriatic keratinocytes is a pivotal molecular event that transduces IL-17 signaling in a keratinocyte-intrinsic fashion. By generating a psoriasis-predisposed mouse model, we were able to elucidate this previously neglected axis.

ARG1 is an M2 macrophage marker that is well-known for restricting NO generation. An intriguing study on patients with psoriasis showed that an NO-donating gel significantly mitigated psoriatic plaques (Abeyakirthi et al., 2010). This finding supports that arginase is overactive in the psoriatic epidermis and infers the potential of metabolic manipulation in psoriasis treatment. Despite this advance, the direct effects of arginase-mediated metabolic fluxes were still ill-defined in both M2 macrophages and in the broader context of immune regulation.

Cellular metabolites can have a profound influence on physiological and pathological processes through their interactions with big molecules, or more specifically, through protein structure modifications (Li et al., 2010). For example, Succinate induces IL-1β secretion by stabilizing HIF-1α in macrophages (Tannahill et al., 2013) and L-Arginine supports T-cell survival by interacting with three nuclear proteins (Geiger et al., 2016). Here, we reveal the functional significance of another aspect of metabolite–big molecule interactions by studying nucleic acids in complex with metabolites. In the context of chronic inflammation, we show these interactions affect the intercellular crosstalk between psoriatic keratinocytes and DCs. Given the central role of nucleic acid sensing in the pathogenesis of various auto-inflammatory diseases, our findings extend our understanding of keratinocyte-featured metabolic scenario and a pro-inflammatory role of keratinocyte-derived metabolites to activate DCs in psoriasis.

Reduced Arginine availability results in T-cell anergy or death (Geiger et al., 2016); we were thus interested in the effects of Arginine consumption by psoriatic keratinocytes on T-cell activity. In this study, K5. Pp6^fl/fl^ mice showed equivalent Arginine levels in the epidermis when compared to littermate controls. Similarly, we found no difference in the serum Arginine abundance between patients with psoriasis and healthy individuals. scRNAseq analysis, however, found that basal keratinocytes from the psoriatic epidermis highly expressed ASS1, the key enzyme required for *de novo* Arginine generation. We assume, therefore, that basal keratinocytes act as a source of cutaneous Arginine, which is the “master and commander” of innate immune responses (Morris, 2010).

To assess the contributions of metabolic rewiring to psoriasis pathology, we used small molecules, including Oligomycin, DFMO and nor-NOHA, to target different metabolic nodes identified from K5. Pp6^fl/fl^ keratinocytes to treat IMQ-induced psoriasis-like skin inflammation. From the perspective of disease mechanism delineation, the validity of these molecules demonstrates the involvement of OXPHOS, polyamine biosynthesis and arginase as pathogenic factors in psoriasis-like inflammation. From the metabolic network perspective, we suggest that increased arginase activity in the psoriatic epidermis is a causative factor for inordinate polyamine generation and decreased NO levels, the latter directly affecting OXPHOS capacity. Phase 1 and phase 2 clinical trials for coronary artery disease (Kovamees et al., 2014) have shown that nor-NOHA is safe and efficient in human patients at both restraining polyamine synthesis and restoring NO production (Li et al., 2016; Moon et al., 2014). We propose that nor-NOHA is a promising drug candidate for psoriasis treatment as its specific target arginase is central in the metabolic adaptation of psoriatic keratinocytes.

In summary, our new mouse model of psoriasis with keratinocyte metabolic rewiring has helped us elucidate the metabolite-facilitated activation of innate immunity. This study presents a previously unrecognized scenario based on self-RNA sensing in chronic skin inflammation. Going forward, a mouse model with impaired polyamine-producing activity in keratinocytes should be generated to investigate the precise contribution of polyamines to self-RNA sensing in psoriasis. Further pre-clinical studies on the therapeutic efficacy of nor-NOHA delivered orally or intravenously to treat psoriasis are needed, so as contrastive researches on nor-NOHA tested together with other anti-psoriasis drugs.

## METHODS

### Human subjects

Psoriatic skin samples were obtained by punch biopsy from patients who were under local lidocaine anesthesia. Normal adult human skin specimens were taken from healthy donors who were undergoing plastic surgery. Serum samples were collected from patients diagnosed with psoriasis vulgaris and healthy donors. Human peripheral blood mononuclear cells (PBMCs) were obtained from blood buffy coats of healthy donors. Patient information is provided in Supplementary Table 1. All participants provided written informed consent. The study was performed in accordance with the principles of the Declaration of Helsinki and approved by the Research Ethics Board of Xiangya Hospital of Central South University, Shanghai General Hospital, Shanghai Tenth People’s Hospital and Changhai Hospital (No. 201311392 and No. 2018KY239).

### Immunohistochemical and histological analysis

Skin specimens were embedded in paraffin and sectioned at the Histology Core of the Shanghai Institute of Immunology. For immunohistochemical staining, the sections were deparaffinized and washed in phosphate-buffered saline (PBS). Antigen retrieval was performed by heating the sections in 10 mM sodium citrate buffer (pH = 6.0). The sections were washed in PBS after cooling, incubated in 3% hydrogen peroxide for 10 min at room temperature (RT) and then washed again in PBS. The sections were blocked in PBS containing 1% bovine serum albumin (BSA) for 1 h at RT and stained in blocking buffer containing primary antibody (anti-PP6C, Merck Millipore cat. 07-1224, 1:50 dilution; anti-mouse Ki67, Servicebio cat. GB13030-2, 1:100 dilution) overnight at 4℃. On the following day, the sections were warmed to RT for 1 h, washed three times in PBS and stained with an HRP-polymer complex for 20 min, which was followed by incubation with secondary antibody for 20 min. The sections were washed three times in PBS, developed with DAB reagent (Peroxidase Substrate Kit, ZSGB-BIO cat. ZLI-9018) and counterstained with hematoxylin. The sections were washed with tap water and then subsequent washes of increasing ethanol concentration for dehydration. Once mounted and air-dried, the slices were viewed under a Zeiss Axio Scope.A1 light microscope equipped with an AxioCam MRc digital camera and analyzed with Axiovision software. Epidermal hyperplasia (acanthosis) was assessed as a histological feature. The pixel size of the epidermal area was measured using the lasso tool in Adobe Photoshop CS4. The absolute area of the epidermis (100 times magnification) was calculated using the following formula: area (mm^2^) = 1.01^2^ × pixels/10^6^. To quantify the expression levels of PP6, the histochemistry score (H score) was applied to evaluate the positive cases, in which both the intensity and the percentage of positivity were considered using the following formula: H score = 3 × (strong intensity) × % + 2 × (moderate intensity) × % + 1 × (mild intensity) × %.

### Animals

Pp6^fl/fl^ mice were provided by Dr. Wufan Tao (State Key Laboratory of Genetic Engineering and Institute of Developmental Biology and Molecular Medicine, Fudan University, Shanghai, China) (Ye et al., 2015). Keratin 5-Cre transgenic mice were provided by Dr. Xiao Yang (State Key Laboratory of Proteomics, Genetic Laboratory of Development and Disease, Institute of Biotechnology, Beijing, China) (Mao et al., 2003). Toll-like receptor 7 targeted mutation (Tlr7^-^) mice were obtained from The Jackson Laboratory (Lund et al., 2004). IL17-GFP mice were provided by Dr. Zhinan Yin (The First Affiliated Hospital, Biomedical Translational Research Institute and Guangdong Province Key Laboratory of Molecular Immunology and Antibody Engineering, Jinan University) (MGI: J:184819). The mice were bred and maintained under specific pathogen-free (SPF) conditions. Age-matched and sex-matched mice were used for all of the experiments in accordance with the National Institutes of Health Guide for the Care and Use of Laboratory Animals with the approval (SYXK-2003-0026) of the Scientific Investigation Board of Shanghai Jiao Tong University School of Medicine in Shanghai, China. To ameliorate any suffering that the mice observed throughout these experimental studies, the mice were euthanized by CO2 inhalation. Healthy Cynomolgus monkeys, ranging in age from 3 to 5 years, were raised in the monkey breeding base of Changchun Biotechnology Development Co., Ltd. (SYXK Gui 2015-0001), Guangxi, China. The managing protocols of the monkeys were carried out in accordance with the standard procedures of Shanghai Hekai Biotechnology Co., Ltd., referring to the Guide for Care and Use of Laboratory Animal (2010) and the principles on Animal Welfare Management (Public Law 99-198).

### Animal models of psoriasis and treatment

For the IMQ-induced mouse model of psoriasis, male C57BL/6 mice (7 weeks-of-age) purchased from Shanghai SLAC Laboratory Animal Co. (Shanghai, China) were maintained under SPF conditions. The mice were subjected to a daily topical dose of 62.5 mg IMQ cream (5%) (MedShine cat. 120503) on the shaved back or 20 mg per ear for six consecutive days. On the indicated days, the skin thickness was measured with a micrometer (Schieblehre). The control mice were treated with an equivalent dose of vehicle cream. For the metabolite treatment, L-Arginine (Sangon Biotech cat. A600205, 5 mg/ml), L-Ornithine (Sigma-Aldrich cat. O6503, 4 mg/ml), L-Proline (Sangon Biotech cat. A600923, 4 mg/ml), Creatine (Sangon Biotech cat. A600326, 10 mg/ml), L-Citrulline (Sangon Biotech cat. A604057, 5 mg/ml) and Spermidine (Sigma-Aldrich cat. S0266, 2 mg/ml) were added to the drinking water 7 days prior to the IMQ application and the treatment lasted until the end of model establishment. For the therapeutic experiments, 25 μl PBS containing DFMO (Sigma-Aldrich cat. D193, 4 μg/μl), nor-NOHA (Cayman cat. 1140844-63-8, 4 μg/μl) and Oligomycin (Abcam cat. 141829, 0.2 μg/μl) were intradermally injected to the dorsal side of the mouse ears daily prior to the IMQ treatment. For the IL-23-mediated mouse model of psoriasis, the ears of the mice were intradermally injected with 500 ng recombinant mouse IL-23 (R&D Systems cat. 1887-ML-010) dissolved in 25 μl PBS into one ear and 25 μl PBS into the contralateral ear. The injections were continued every other day for a total of eight injections. The ears were collected for analysis on day 16. To treat K5. Pp6^fl/fl^ mice, an intraperitoneal injection of 100 μl PBS containing nor-NOHA (2 μg/μl) or DFMO (1 μg/μl) or Oligomycin (0.1 μg/μl) was performed every other day for 2 weeks. All procedures were approved and supervised by Shanghai Jiao Tong University School of Medicine Animal Care and Use Committee. For the IMQ-induced non-human primate model of psoriasis, the Cynomolgus monkeys were depilated on the back shoulders and subjected to a daily dose of 120 mg IMQ cream in the morning for 10 days. An intradermal injection of 50 μl PBS containing nor-NOHA (10 μg/μl) was performed every afternoon post-IMQ application. An equal volume of PBS was injected to the contralateral shoulder of the monkey. The clinical scores of the skin phenotype were graded according to Lattice System Physician’s Global Assessment (Langley and Ellis, 2004).

### Immunoprecipitation and immunoblotting

For PP6 immunoprecipitation, nuclear and cytoplasmic extracts from HaCaT cells were prepared with a Nuclear Protein Extraction Kit (Beyotime cat. P0027) supplemented with a protease inhibitor cocktail (Bimake cat. B14001). The lysates were incubated with anti-PPP6C (Abcam cat. 131335, 1:50) or rabbit IgG (Abcam cat. 172730) overnight at 4℃, followed by binding of protein A/G magnetic beads (Merck Millipore cat. LSKMAGAG02) for 4 h at 4℃. The beads were rinsed three times with a wash buffer and then eluted with SDS loading buffer. The immunoprecipitates were then subjected to immunoblotting. For Spermidine/Spermine immunoprecipitation, anti-Spermine (Abcam cat. 26975) or rabbit IgG was cross-linked to protein A/G magnetic beads (Bimake cat. B23202) using dimethyl pimelimidate (ThermoFisher Scientific cat. 21667). Plasma samples from psoriasis patients and healthy donors were pre-cleared with rabbit IgG and protein A/G magnetic beads, and then incubated with the antibody-cross-linked beads overnight at 4℃. The beads were rinsed three times with a wash buffer and eluted with 50 mM Glycine (pH = 2.8). The immunoprecipitates were neutralized with 1 M Tris-HCl (pH = 7.5) and subjected to protein mass spectrometry. For immunoblotting, mouse skins, cultured cells or nuclear fractions were lysed in radioimmunoprecipitation assay buffer containing a protease and phosphatase inhibitor cocktail (ThermoFisher Scientific cat. 78440). Anti-PPP6C (Abcam cat. 131335, 1:1000 dilution), anti-β-actin (Proteintech cat. 60008-1-Ig, 1:20000 dilution), anti-C/EBPβ (Santa Cruz cat. 7962, 1:1000 dilution), anti-p-C/EBPβ (Thr188) (Cell Signaling Technology cat. 3084, 1:1000 dilution), anti-ARG1 (Proteintech cat. 66129-1-Ig, 1:1000 dilution), anti-SDHA (Proteintech cat. 14865-1-AP, 1:1000 dilution), anti-MTCO2 (Proteintech cat. 55070-1-AP, 1:1000 dilution), anti-COX4I1 (Proteintech cat. 11242-1-AP, 1:1000 dilution), anti-IkappaB-ζ (Cell Signaling Technology cat. 93726, 1:1000 dilution), anti-alpha Tubulin (Proteintech cat. 66031-1-Ig, 1:2000 dilution), anti-GAPDH (Proteintech cat. 60004-1-Ig, 1:10000 dilution), anti-Histone H3 (Cell Signaling Technology cat. 4620, 1:1000 dilution), HRP-labeled goat anti-mouse IgG (H+L) (Beyotime cat. A0216, 1:1000 dilution) and HRP-labeled goat anti-rabbit IgG (H+L) (Beyotime cat. A0208, 1:1000 dilution) were used as the antibodies. The signal was detected with ECL Western Blotting Substrate (ThermoFisher Scientific cat. 34095) and an Amersham Imager 600 (GE Healthcare). The images were cropped for presentation.

### Flow cytometry

Single cell suspensions generated from the skin of Pp6^fl/fl^, K5. Pp6^fl/fl^ or C57BL/6 (IMQ treated or not) mice were stained with fluorophore-conjugated antibodies. After washing, the cells were assayed with a BD LSRFORTESSA flow cytometer, and the data were analyzed using FlowJo software. Monoclonal antibodies against CD45 (clone 30-F11), CD3 (clone 145-2C11), CD11c (clone N418), F4/80 (clone BM8), Ly6G (clone 1A8-Ly6g), CD31 (clone PECAM-1), CD80 (clone 16-10A1), CD86 (clone GL1), MHC Class II (clone M5/114.15.2) which were obtained from eBioscience, were used in the flow cytometry analyses.

### Immunofluorescence

Cryosections (10 1m) of mouse or monkey skin were cut using a cryostat (Leica, CM1950) and fixed in ice-cold acetone for 15 min. Human paraffin sections were dewaxed and antigen unmasked. The slices were blocked with 1% BSA/PBS for 1 h and stained with anti-Krt5 (Invitrogen cat. MA5-17057, 1:100 dilution), anti-PPP6C (Abcam cat. 131335, 1:50 dilution), anti-mouse Ki67 (Servicebio cat. GB13030-2, 1:50 dilution), anti-Krt6 (Proteintech cat. 10590-1-AP, 1:2000 dilution), anti-mouse CD80 (BioLegend cat. 104702, 1:50 dilution), anti-eosinophil peroxidase (EPX, Santa Cruz cat. 19148, 1:50 dilution), anti-C/EBPβ (Santa Cruz cat. 7962, 1:100 dilution), anti-p-C/EBPβ (Thr235) (Cell Signaling Technology cat. 3084, 1:100 dilution), anti-ARG1 (Proteintech cat. 66129-1-Ig, 1:1000 dilution), anti-COX4I1 (Proteintech cat. 11242-1-AP, 1:100 dilution), anti-ASS1 (Proteintech cat. 16210-1-AP, 1:100 dilution), anti-human CD4 (BioLegend cat. 317402, 1:50 dilution) and anti-human Integrin α-X (Santa Cruz cat. 1185, 1:50 dilution) overnight at 4℃. Thereafter, the slices were rinsed three times in PBS and stained with fluorochrome-conjugated secondary antibodies (all from Life Technologies) at a 1:500 dilution for 1 h at RT in the dark. DAPI (BD Biosciences cat. 564907) incubation at a 1:2,000 dilution at RT for 5 min was used for nuclei staining. After being washed in PBS, the slices were mounted with a fluorescent mounting medium (Dako cat. S3023) and visualized under a confocal microscope (Leica, TCS SP8).

### TEWL measurement

TEWL measurements were carried out at RT and at a humidity level of 50 ± 20%. The experiments were performed in triplicate for each spot using a gpskin™ Barrier Light skin barrier function measurement device.

### Cell culture

To isolate primary murine keratinocytes, the skin from the tails of adult mice was cut into small pieces and then suspended on Dispase II (Sigma-Aldrich cat. D4693, 2 mg/ml) overnight at 4℃. The following day, the epidermis was peeled and placed into 0.05% trypsin-EDTA (Gibco cat. 25300062) for 10 min at 37℃ with gentle shaking. The cell suspension was added to a cold trypsin neutralization solution and then centrifuged at 400 g for 10 min. The cells were cultured in Medium 154CF (Gibco cat. M154CF500) supplemented with 0.05 mM calcium chloride and human keratinocyte growth supplement (Gibco cat. S0015). HaCaT cells and NIH3T3 cells were cultured in DMEM/high glucose (HyClone cat. SH30022.01) containing 10% fetal bovine serum (FBS). Normal Human Epidermal Keratinocytes (NHEKs; Lifeline Cell Technology, cat. FC-0007) stored in liquid nitrogen were defrosted and cultured in serum-free basal medium with growth factors (Lifeline Cell Technology, cat. LL-0007). The medium was refreshed every 2 days and the cells were sub-cultured according to the cell fusion. Cells at passage 2-6 were used for subsequent experiments. Immature BMDCs were harvested from femurs and tibias of 8-to12-week-old mice and cultured in the presence of recombinant mouse GM-CSF (R&D Systems cat. 415-ML, 20 ng/ml) and IL-4 (R&D Systems cat. 404-ML, 10 ng/ml) in RPMI 1640 medium (Gibco cat. 11875093) supplemented with 10% FBS and 2-Mercaptoethanol (Gibco cat. 21985023) for 6 days. Immature human moDCs were generated from PBMCs using CD14 Microbeads (Miltenyi Biotec cat. 130-050-201) and cultured for 6 days with recombinant GM-CSF (R&D Systems cat. 215-GM, 50 ng/ml) and IL-4 (R&D Systems cat. 204-IL, 35 ng/ml) in RPMI 1640 medium supplemented with 10% FBS and 2-Mercaptoethanol. Mouse pDCs were generated in the presence of 200 ng/mL recombinant Flt3l (PeproTech cat. 250-31L) from bone marrow cells and purified by fluorescence-activated cell sorting (FACS) for B220^+^CD317^+^ cells.

### RNA-seq and qPCR

Skin biopsy specimens were snap-frozen in liquid nitrogen and pulverized. HaCaT cells were rinsed with PBS. Total RNA was isolated using TRIzol Reagent (Invitrogen cat. 15596026) and quantified with a NanoDrop spectrophotometer. For RNA-seq, a cDNA library was prepared using a TruSeq RNA sample preparation kit (Illumina cat. FC-122-1001) according to the manufacturer’s instructions. The samples were quantified using a TBS-380 microfluorometer with Picogreen reagent, clustered with a TruSeq paired-end cluster kit v3-cBot-HS (Illumina cat. PE-401-3001) and sequenced on an Illumina HiSeq2000 platform (100 bp, TruSeq SBS kit v3-HS, 200 cycles). The adaptor sequences were trimmed from the raw paired-end reads, and RNA *de novo* assembly was achieved using TopHat (http://tophat.cbcb.umd.edu/). The read counts were further normalized into FPKM values, and the fold changes were calculated using Cufflinks (http://cole-trapnell-lab.github.io/cufflinks/). DEGs were identified using Cuffdiff (http://cole-trapnell-lab.github.io/cufflinks/cuffdiff/). Cross-species analyses were joined by case-insensitive gene symbol matching. For the psoriasis analysis, the RNA-seq data set from 82 normal skin samples and 92 psoriasis skin samples (accession no. GSE54456) was used. For the atopic dermatitis analysis, 30 human skin samples (15 samples from subjects with atopic dermatitis and 15 samples from healthy subjects) were obtained from arrays (accession no. GSE12511). Pathway enrichment analysis was performed using the clusterProfiler package (DOI: 10.18129/B9.bioc.clusterProfiler).

For qPCR detection of *ARG1*, total RNA from HaCaT cells was reverse-transcribed into cDNA using an M-MLV First-Strand Synthesis Kit (Invitrogen cat. C28025-021) with oligo(dT) primers. qPCR was conducted with the FastStart Universal SYBR Green Master (Roche cat. 04913914001) in a ViiA 7 Real-Time PCR system (Applied Biosystems). The relative expression of target genes was confirmed using the quantity of target gene/quantity of *GAPDH*. For qPCR detection of human 28S rRNA, endosomes from mouse BMDCs treated with RNA complexes for 4 h were collected using a Minute™ Endosome Isolation and Cell Fractionation Kit (Invent Biotech cat. ED-028) and endosomal RNA was extracted and then reversed-transcribed with random primers. The following primers were used: *GAPDH* forward:

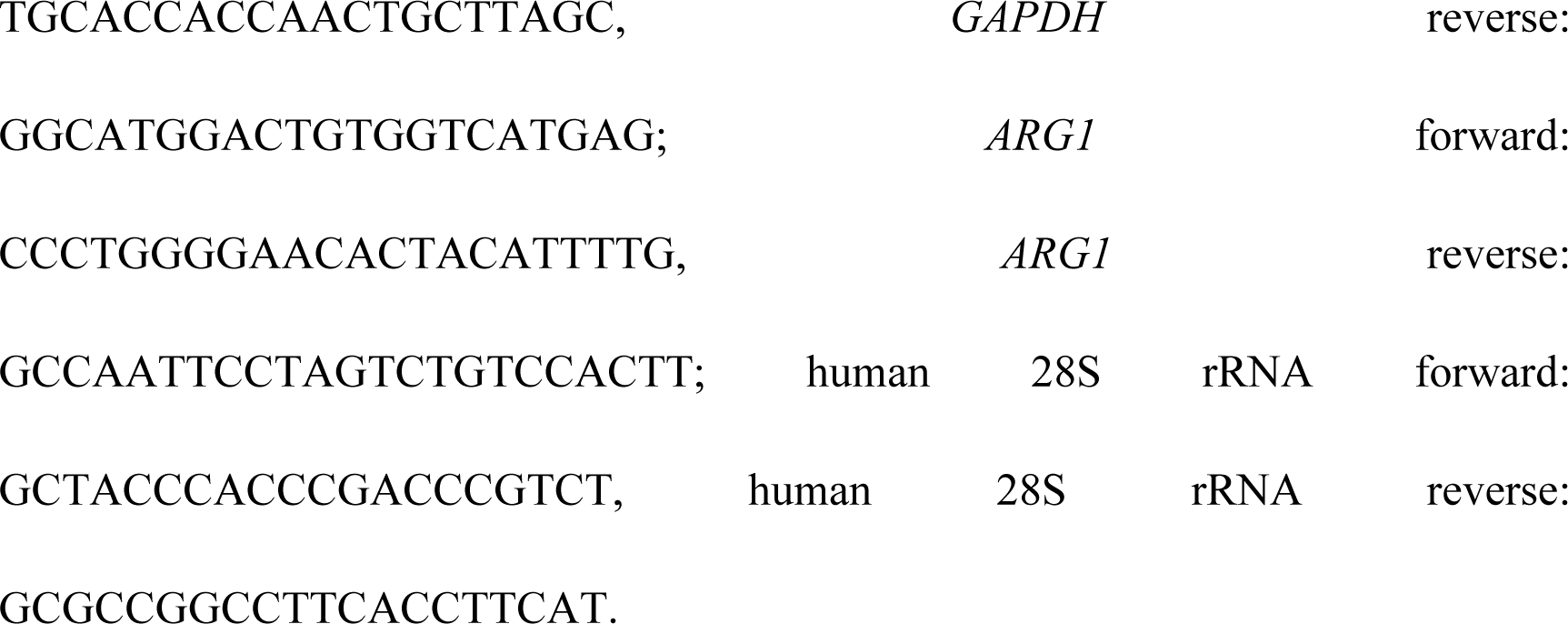

### Semi-quantitative cytokine array

The cytokine levels in mouse skin were measured with a Mouse Cytokine Array GS4000 (RayBiotech cat. GSM-CAA-4000-1) according to the manufacturer’s instructions. The fluorescein-labeled array was visualized using an InnoScan 300 Microarray Scanner. Data were extracted by GenePix Pro 5.1 software and analyzed with RayBiotech Q-Analyzer software.

### ELISA

The serum IgE levels of the mice were measured using Mouse IgE ELISA MAX™ Deluxe (BioLegend cat. 432404) according to the manufacturer’s instructions. To detect murine IL-6 and Tnf in BMDC cultures, a Mouse IL-6 ELISA kit (NeoBioscience cat. EMC004.96) and a Mouse TNF-α ELISA kit (NeoBioscience cat. EMC102a.96) was used. To detect human IL-6 and TNF-α in moDC cultures, a Human IL-6 ELISA Kit (MultiSciences cat. 70-EK106/2-96) and a Human TNF-α High Sensitivity ELISA Kit (MultiSciences cat. 70-EK182HS-96) was used. To detect murine Ifn-α in pDC cultures, a Mouse IFN-alpha ELISA Kit (R&D Systems cat. 42120-1) was used.

### Halometasone painting and IL-17A neutralization treatment

Halometasone cream (50 mg; Bright Future Pharmaceutical Laboratories Limited) was topically painted onto the plaques of affected K5. Pp6^fl/fl^ mice every other day for 2 weeks. Then, 100 μg anti-mouse IL-17A (BioxCell cat. BE0173) or isotype control (BioxCell cat. BE0083) was intraperitoneally injected into the affected K5. Pp6^fl/fl^ mice once each week for a month.

### RNA interference

Small interfering RNA (siRNA) targeting human PPP6C was synthesized by GenePharma: sense: CUAAAUGGCCUGAUCGUAU, anti-sense: AUACGAUCAGGCCAUUUAG. The siRNA molecules were transfected using a TransIT-X2 Dynamic Delivery System (Mirus cat. MIR 6000).

### ChIP-qPCR

ChIP was performed using the SimpleCHIP enzymatic chromatin immunoprecipitation kit (Cell Signaling Technology cat. 9002) according to the manufacturer’s instruction with minor modifications. In brief, primary mouse keratinocytes were harvested and cross-linked with 1% (v/v) formaldehyde for 10 min at RT. Subsequently, the nuclei were isolated by lysing the cytoplasmic fraction, and chromatin was digested into fragments of 150–900 bp with micrococcal nuclease for 20 min at 37℃, followed by ultrasonic disruption of the nuclear membrane using a standard microtip and a Branson W250D Sonifier (four pulses, 60% amplitude and duty cycle 40%). The sonicated nuclear fractions were divided for input control and for overnight incubation at 4℃ with 10 μg of either anti-C/EBPβ (Abcam cat. 32358) or the negative control IgG. After incubation with 30 μl ChIP grade protein G-agarose beads for 2 h at 4 °C, the Ab-protein-DNA complexes were eluted from the beads and digested by proteinase K (40 μg) for 2 h at 65 ℃, followed by spin column-based purification of DNA. Finally, genomic DNA recovered from the ChIP were qPCR amplified with primers specific to the C/EBPβ binding elements of the *Arg1* promoter region. The primers used to detect the *Arg1* promoter sequence were as follows: forward: CCCAAAGTGGCACAACTCAC, reverse: GCAGAAGGCTTTGTCAGCAG. The signals were expressed as a percentage of the total input chromatin.

### ScRNAseq

Fresh skin was placed in saline at 4℃ until further processing. Single cell suspensions were generated by enzyme digestion and subjected to FACS to exclude doublets, debris and DAPI-positive dead cells. Sorted cells were centrifuged and resuspended in 0.04% BSA in PBS. Chromium Single Cell 3’ v3 (10X Genomics) library preparation was conducted by the Sequencing Core at the Shanghai Institute of Immunology, according to the manufacturer’s instructions. The resulting libraries were sequenced with an Illumina HiSeq 4000 platform. Raw data were processed using Cell Ranger (version 3.0) and further filtered, processed and analyzed using the Seurat package (Satija et al., 2015).

### Proximity Ligation Assay (PLA)

HaCaT cells were seeded on glass coverslips and transfected with plasmids overexpressing WT C/EBP-β, C/EBP-β^S184A^, C/EBP-β^T188A^, and double mutant C/EBP-β for 24 h. The cells were fixed and permeabilized with 4% PFA in PBS for 15 min at RT. PLA was performed using Duolink® In Situ Red Starter Kit Mouse/Rabbit (Sigma-Aldrich cat. DUO92101-1KT) according to manufacturer’s instructions. Briefly, blocking was performed with blocking solution for 30 min at 37℃, and primary antibodies against PP6 (Abcam cat. 131335, 1:50 dilution) and Flag-tag (MBL cat. M185-3L, 1:10000 dilution) were incubated overnight at 4℃. The PLA Probe anti-mouse PLUS and PLA Probe anti-rabbit MINUS were incubated for 1 h at 37℃. Ligation and amplification were performed using Detection Reagents Red. Duolink® Mounting Medium with DAPI was used for nuclear staining and mounting.

### NO quantification

Intracellular NO levels were quantified using an OxiSelect™ Intracellular Nitric Oxide (NO) Assay Kit (Cell Biolabs cat. STA-800). Primary mouse keratinocytes were incubated with medium containing Nitric Oxide Fluorometric Probe at a 1:1000 dilution at 37℃ for 30 min. The cells were rinsed, digested and analyzed on a BD FACSCanto II cell analyzer using FITC filter set.

### Extracellular flux analysis

Oxygen consumption rate (OCR) and extracellular acidification rate (ECAR) measurements were performed using an XFe96 Extracellular Flux analyzer (Seahorse Bioscience) according to the manufacturer’s instructions. Briefly, keratinocytes were seeded into an XFe96 polystyrene cell culture plate at a density of 5 × 10^4^ cells/well and allowed for attachment overnight. Prior to the assay, the growth medium was exchanged with the appropriate assay medium, and the cells were incubated in a 37℃/non-CO2 incubator for 1 h. Three baseline measurements were taken, and three response measurements were taken after the addition of an indicated compound. OCR and ECAR were reported as absolute rates (pmoles/min for OCR and mpH/min for ECAR), normalized against the protein concentrations measured by BCA method. For the mitochondrial fuel flexibility assessment, Seahorse XF Mito Fuel Flex Test Kit (Agilent Technologies cat. 103260-100) was used according to the manufacturer’s instructions. The results were calculated according to the formulas: Dependency% = ((Baseline OCR – Target inhibitor OCR) / (Baseline OCR – All inhibitors OCR)) × 100; Capacity% = (1 – (Baseline OCR – Other inhibitors OCR) / (Baseline OCR – All inhibitors OCR) × 100; Flexibility% = Capacity% – Dependency%.

### ATP measurement

The keratinocyte ATP content was measured using the ENLITEN® ATP Assay System (Promega cat. FF2000). Primary mouse keratinocytes were seeded at a density of 10^4^ cells/well in a 96-well plate and allowed for attachment overnight. The cells were lysed with 20 μl 0.5% trichloroacetic acid and shaken for 5 min at RT before neutralization with 80 μl Tris-acetate (pH = 7.8). The extract was mixed with rL/L Reagent and the ATP level was measured using a luminescent plate reader (BioTek).

A standard curve of ATP was made, and the protein concentrations were measured by BCA method using extra cells.

### Unbiased metabolomics analysis

Epidermis samples from the tail skin of eight Pp6^fl/fl^ and twelve K5. Pp6^fl/fl^ mice were collected. The samples were snap frozen and metabolites were extracted with 80% ice-cold methanol. Liquid chromatography-mass spectrometry (LC-MS) was performed on an Ultimate 3000-Velos Pro system equipped with a binary solvent delivery manager and a sample manager, coupled with an LTQ Orbitrap Mass Spectrometer equipped with an electrospray interface (ThermoFisher Scientific). The raw data were processed with progenesis QI (Waters Corporation) and Principal component analysis (PCA) and hierarchical clustering of metabolites were analyzed.

### Targeted metabolite quantitation

The sera were loaded onto an ACQUITY UPLC HSS T3 Column (2.1 × 100 mm, 1.8 μm, Waters Corporation), and separated with mobile phase A (10 mM NH4COOH) and mobile phase B (100% acetonitrile) at a flow rate of 0.3 ml/min with column temperature set at 40℃. Monitoring of the analytes and their respective internal standards was achieved using an UltiMate 3000 UHPLC/TSQ Vantage LC-MS/MS system (ThermoFisher Scientific) equipped with a heated electrospray ionization source (HESI). The instrument was operated in selected reaction monitoring (SRM) mode. The mass analyzer settings were as follows: Vaporizer temperature, 300°C; spray voltage, positive polarity 3000 V/negative polarity 2500 V; capillary temperature, 300 °C; sheath gas pressure, 40; aux gas pressure, 10; ion sweep gas pressure, 0; collision gas pressure (mTorr), 1.5; Q1 Peak width (FWHM), 0.7; Q3 Peak width (FWHM), 0.7; cycle time (s), 0.4 s.

### Protein mass spectrometry

The immunoprecipitates pulled down with anti-Spermidine/Spermine or Rabbit IgG from human plasma samples and sera samples from mice were filtered with Amicon® Ultra Centrifugal Filters to segregate ingredients ≤ 3 kD or > 3 kD. Samples ≤ 3 kD were left untreated until lyophilization. Samples > 3 kD were dissolved in a denaturation buffer (8 M Urea, 0.1 M Tris-HCl, pH = 8.5) and centrifuged at 14,000 g for 20 min at 20℃ in YM-10 filter units three times. Mixed protein lysates were reduced with 10 mM dithiothreitol (DTT) for 1 h at RT and then alkylated with 55 mM iodoacetamide (IAA) for 1 h at RT in the dark. The urea solvent of the protein mixtures was exchanged with 50 mM NH4HCO3 by centrifugation at 14,000 g for 20 min at 20℃ three times, followed by protein digestion with sequencing grade modified trypsin (Promega, cat. V5111) at a protein-to-enzyme ratio of 50:1 at 37℃ overnight. Tryptic peptides were collected by centrifugation at 14,000 g for 20 min at 20 ℃. The Tryptic peptides were treated with 1% trifluoroacetic acid (TFA) and purified using the C18 Ziptips and eluted with 0.1% TFA in 50∼70% acetonitrile. The eluted peptides and the untreated samples ≤ 3 kD were lyophilized using a SpeedVac (ThermoSavant) and resuspended in 10 μl 1% formic acid/5% acetonitrile. All mass spectrometric experiments are performed on a QE-Plus mass spectrometer connected to an Easy-nLC 2000 via an Easy Spray (ThermoFisher Scientific). The peptide mixtures were loaded onto a 15 cm column with 0.075 mm inner diameter packed with C18 2-μm Reversed Phase resins (PepMap RSLC), and separated within a 60 min-linear gradient from 95% solvent A (0.1% formic acid / 2% acetonitrile / 98% water) to 28% solvent B (0.1% formic acid / 80% acetonitrile) at a flow rate of 300 nl/min. The spray voltage was set to 2.1 KV, with the temperature of ion transfer capillary set at 275℃, and the RF lens was 60%. The mass spectrometer was operated in positive ion mode and employed in the data-dependent mode to automatically switch between MS and MS/MS using the Tune and Xcalibur 4.0.27.19 software package. One full MS scan from 350 to 1,500 m/z was acquired at high resolution R = 70,000 (defined at m/z=400), followed by fragmentation of the 20 most abundant multiply charged ions (singly charged ions and ions with unassigned charge states were excluded), for ions with charge states 2–7 and collision energy of 27% ± 5%. Dynamic exclusion was used automatically. All MS/MS ion spectra were analyzed using PEAKS 8.0 (Bioinformatics Solutions) for processing, *de novo* sequencing and database searching. The resulting sequences were searched through the UniProt Human Proteome database (downloaded on May 5^th^, 2018) with the mass error tolerance set at 10 ppm and 0.02 Da for parent and fragment, respectively. FDR estimation was enabled. Peptides were filtered for −log10P ≥ 20, and the proteins were filtered for −log10P ≥ 15 plus one unique peptide. For all experiments, these settings gave an FDR of < 1% at the peptide-spectrum match level.

### DC stimulation

Synthetic HNRNPA1226-237 and Tetramethylrhodamine (TMR)-labeled HNRNPA1226-237 (purity > 95%) were purchased from GeneScript. Mouse BMDCs were seeded in 96-well plates at a density of 10^5^/well. Total mouse RNA (self-RNA) was extracted from NIH3T3 cell line. Self-RNA (5 μg/ml as a final concentration) was premixed with the indicated reagents (all at 1 μg/ml as final concentrations) in RPMI 1640 medium for 20 min at RT and then added to the BMDC cultures. Human moDCs were seeded in 96-well plates at a density of 5 × 10^4^/well. Total human RNA (self-RNA) was extracted from HaCaT cell line. Self-RNA (10 μg/ml as a final concentration) was premixed with the indicated reagents (all at 2 μg/ml as final concentrations) in RPMI 1640 medium for 20 min at RT and then added to the moDC cultures. Mouse pDCs were seeded in 96-well plates at a density of 5 × 10^4^/well. Mouse DNA (self-DNA) was extracted from NIH3T3 cell line. Self-DNA (5 μg/ml as a final concentration) was premixed with the indicated reagents (all at 1 μg/ml as final concentrations) in RPMI 1640 medium for 20 min at RT and then added to the pDC cultures. For ELISA, the supernatants were collected after overnight culture. For microscopic imaging, self-RNA was labeled using a Label IT® Nucleic Acid Labeling Kit (Mirus cat. MIR7125), and Spermidine was labeled with 2-(methylamino) benzoic acid (Wolf et al., 2006). BMDCs were collected after 4 h-incubation, fixed with 4% paraformaldehyde and stained with anti-MHC Class II (eBioscience cat. 47-5321-80). For flow cytometric analysis of RNA internalization, BMDCs were treated with 488-labeled self-RNA for 4 h. For flow cytometric analysis of DC activation markers, DCs were isolated from spleens and lymph nodes of C57BL/6 mice using a MojoSort™ Mouse Pan Dendritic Cell Isolation Kit (BioLegend cat. 480097). DCs were cultured with supernatants of necrotic keratinocytes generated by an overnight culture of UV-irradiated (2 J/m^2^/s for 40 min) keratinocytes derived from Pp6^fl/fl^ and K5. Pp6^fl/fl^ mice. After overnight culture, CD80, CD86 and MHC class II levels of CD11c^+^ DCs were detected.

### RNase protection assay

Suspensions containing indicated self-RNA-polyamine-Hnrnpa1226-237 complexes were incubated with Ribonuclease A (Sigma-Aldrich cat. R6513, 1 ng/ml) at 37°C for 5 min and then run on a 1% agarose gel.

### DC and T-cell co-culture

BMDCs were treated with different combinations of self-RNA, polyamines and Hnrnpa1226-237 for 3 h. In the meantime, naïve CD4^+^ T cells were isolated from the spleens of C57BL/6 mice or IL17-GFP mice using a MojoSort™ Mouse CD4 Naïve T Cell Isolation Kit (BioLegend cat. 480040). T cells from C57BL/6 mice were further labeled with CellTrace™ Violet (Invitrogen cat. C34557). BMDCs were washed three times and co-cultured with naïve CD4^+^ T cells at a ratio of 1:10 for 72 h. To detect IL17-GFP, cells were stimulated with 750 ng/ml ionomycin (Sigma-Aldrich cat. I9657), 50 ng/ml phorbol 12-myristate 13-acetate (PMA) (Sigma-Aldrich cat. 79346) and GolgiPlug (BD Biosciences cat. 555029) for another 4 h at 37 ℃. The cells were then collected, stained with antibodies against CD4 (eBioscience cat. 17-0041-82) and CD44 (eBioscience cat. 12-0441-82) and analyzed on a BD FACSCanto II cell analyzer.

### RNAscope combined with immunofluorescence

Cell suspensions from C57BL/6 mouse skin treated with IMQ for 3 days or not were subjected to Percoll™-facilitated gradient separation. DCs were isolated from the resulted mononuclear cells using a MojoSort™ Mouse Pan Dendritic Cell Isolation Kit (BioLegend cat. 480097). Single molecule FISH experiments were carried out using an RNAscope® Multiplex Fluorescent Assay v2 along with an RNAscope® Probe-Mm-Krt14-C2 (ACD cat. 422521-C2). Immunofluorescence was then performed using antibodies against Tlr7 (Novus cat. NBP2-27332, 1:50 dilution) and Spermidine/Spermine (Novus cat. NB100-1847, 1:50 dilution).

### Statistical analysis

The data were analyzed using GraphPad Prism 7 and are presented as the means ± SEM. A Student’s *t*-test was used to compare two conditions, and an analysis of variance (ANOVA) with Bonferroni or Newman-Keuls correction was used for multiple comparisons. A simple linear regression model was used to analyze the correlation between PP6 expression and H score. The probability values of < 0.05 were considered statistically significant, and two-sided Student’s *t*-tests and ANOVA were performed. \**p* < 0.05; \*\**p* < 0.01; \*\*\**p* < 0.001; \*\*\*\**p* < 0.0001; ns: not significant. The error bars depict the SEM.

## Data Availability

The data that support the findings of this study are available from the corresponding author upon reasonable request. RNA-Seq datasets were deposited in GEO under the accession numbers: GSE123019 and GSE122665.

## ACKNOWLEDGMENTS

The authors would like to thank Ms. C. Gao, Ms. M. Zhang, Ms. L. Ding and Dr. L. Chen (Sequencing Core of Shanghai Institute of Immunology) for single-cell cDNA library generation; Dr. L. Xia and Dr. L. Meng (Proteomics Core of College of Basic Medical Sciences, SJTU-SM) for protein LC-MS analyses; Dr. D. Han and Dr. J. Fang (Metabolomics Core of College of Basic Medical Sciences, SJTU-SM) for metabolite quantitation analyses; and Insight Editing London for proofreading the manuscript prior to publication. This work was supported by grants from the National Natural Science Foundation of China (No: 81725018, No: 31330026, No: 31570922 and No: 81803123), the National Program on Key Basic Research Project (973 program, No: 2014CB541905) and the Shanghai Municipal Commission (No: 201740063).

## AUTHOR CONTRIBUTIONS

F.L., Y.S. and HL.W. designed the study and prepared the manuscript. F.L. and Y.S. conducted the experiments and analyzed the data. HL.W. and F.G. supervised the research. Z.X., L.N., Z.W., S.D., Z.L., H.Z., J.B., Q.Y., X.C., L.S. and H.W. helped with experimental details. Z.W., X.C., Y.S. and W.T. reviewed and edited the manuscript.

---

**Figure S1.**
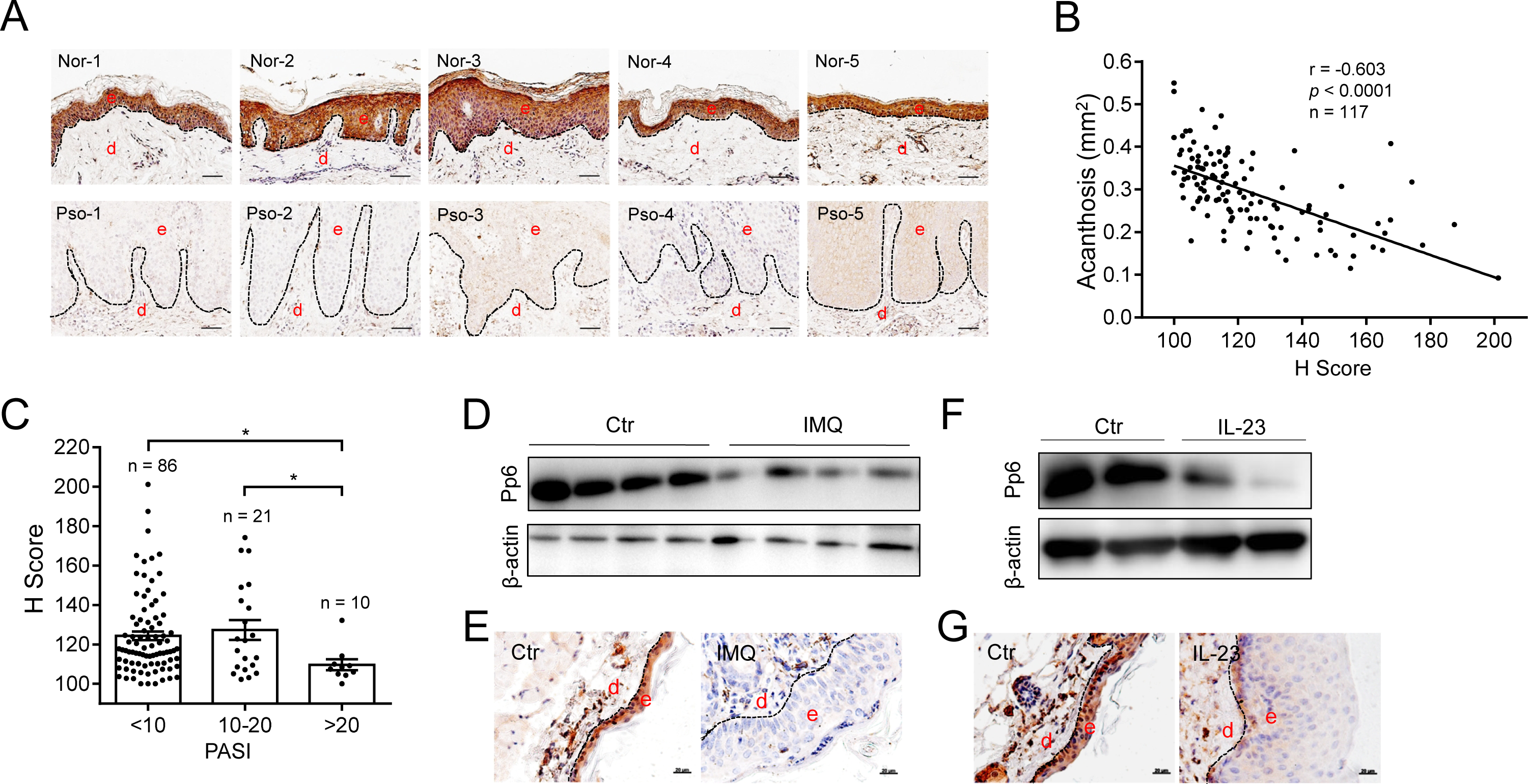
PP6 is diminished in the epidermis of psoriasis and its mouse models. (**A**) Immunohistochemical staining for PP6 in five healthy and five psoriatic human skin sections. Nor, normal skin; Pso, psoriatic skin. (**B**) Linear regression analysis of PP6 expression levels and acanthosis of 117 psoriatic skin biopsies (x axis: PP6 staining score; y axis: acanthosis of the skin section; *r* = -0.603). (**C**) The 117 psoriatic skin samples were separated into three groups according to the patient PASI. The PP6 staining score was compared between these groups. (**D** and **E**) Immunoblotting and immunohistochemical analysis of Pp6 expression in mouse skin derived from mice treated with vehicle (Ctr) or IMQ (n = 4). (**F** and **G**) Immunoblotting and immunohistochemical analysis of Pp6 expression in mouse ears injected with PBS (Ctr) or IL-23 (n = 2). For (**A**, **E** and **G)**, the dotted lines indicate the border between the epidermis and the dermis; e, epidermis; d, dermis; scale bars represent 50 μm (**A**) or 20 μm (**E** and **G**). The data (**D**-**G**) are representative of three independent experiments. \**p* < 0.05, two-tailed Student’s *t*-test (means ± SEM).

**Figure S2.**
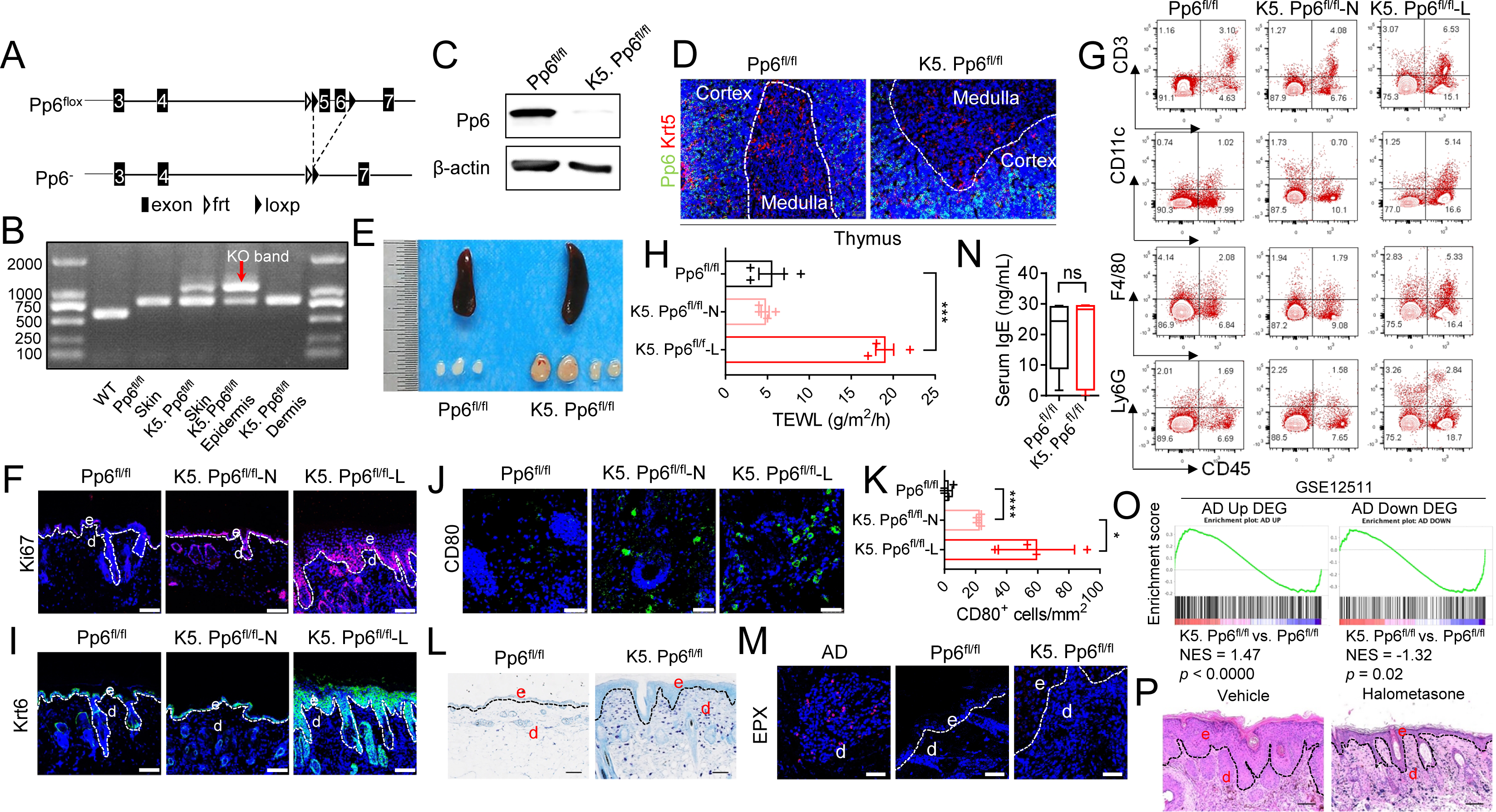
Characterization of skin lesions observed in K5. Pp6^fl/fl^ mice. (**A**) Schematic of *Pp6* deletion in K5. Pp6^fl/fl^ mice. (**B**) *Pp6* genotyping of the indicated tissue samples derived from WT, Pp6^fl/fl^ and K5. Pp6^fl/fl^ mice. The red arrow shows K5-Cre-mediated epidermal-specific deletion of *Pp6*. (**C**) Immunoblot analysis of Pp6 expression in primary murine keratinocytes from Pp6^fl/fl^ and K5. Pp6^fl/fl^ mice. (**D**) Immunofluorescent staining of Krt5 and Pp6 in thymic sections from Pp6^fl/fl^ and K5. Pp6^fl/fl^ mice. The dotted lines indicate the border between the cortex and the medulla; the scale bars represent 50 μm. (**E**) Phenotypic representation of splenomegaly and lymphadenopathy observed from diseased K5. Pp6^fl/fl^ mice in contrast to Pp6^fl/fl^ controls. (**F**) Immunofluorescent labelling of Ki67 in mouse skin samples. (**G**) Flow cytometric analysis of immune cell-specific markers in mouse skin (n = 5∼6). (**H**) TEWL of mouse skin (n = 4). (**I**) Immunofluorescent labelling of Krt6 in mouse skin samples. (**J** and **K**) Immunofluorescent staining and quantitation of dermal CD80^+^ cells of mouse skin samples (n = 4). (**L**) Toluidine blue staining for mast cells in skin sections from Pp6^fl/fl^ and K5. Pp6^fl/fl^ mice. (**M**) Eosinophil peroxidase (EPX) staining for eosinophils in skin sections from Pp6^fl/fl^ and K5. Pp6^fl/fl^ mice. A skin section from a patient diagnosed with atopic dermatitis (AD) served as a positive control. (**N**) Serum IgE levels in Pp6^fl/fl^ and K5. Pp6^fl/fl^ mice, as measured by ELISA (n = 11∼12). Boxes show twenty-fifth to seventy-fifth percentiles; whiskers show minimum to maximum; and lines show medians; ns: not significant, two-tailed Student’s *t*-test. (**O**) GSEA of DEGs in K5. Pp6^fl/fl^ skin relative to Pp6^fl/fl^ skin with upregulated (AD UP) or downregulated (AD DOWN) DEGs in human AD (GSE12511) enriched. (**P**) H&E staining of skin sections from K5. Pp6^fl/fl^ mice treated with Halometasone or vehicle. Pp6^fl/fl^, healthy skin sample from control mice; K5. Pp6^fl/fl^-N, unaffected skin from K5. Pp6^fl/fl^ mice; K5. Pp6^fl/fl^-L, affected skin from K5. Pp6^fl/fl^ mice. The data (**D, E, I, L** and **M**) are representative of > 5 mice in each group. The data (**P**) are representative of two independent experiments with two mice in each group. For (**F**, **I**, **J**, **L**, **M** and **P**), the dotted lines indicate the border between the epidermis and the dermis; e, epidermis; d, dermis; the scale bars represent 50 μm (**F**, **I**, **J**, **M** and **P**) or 100 μm (**L**). \**p* < 0.05, \*\*\**p* < 0.001, \*\*\*\**p* < 0.0001, ns: not significant, two-tailed Student’s *t*-test (means ± SEM).

**Figure S3.**
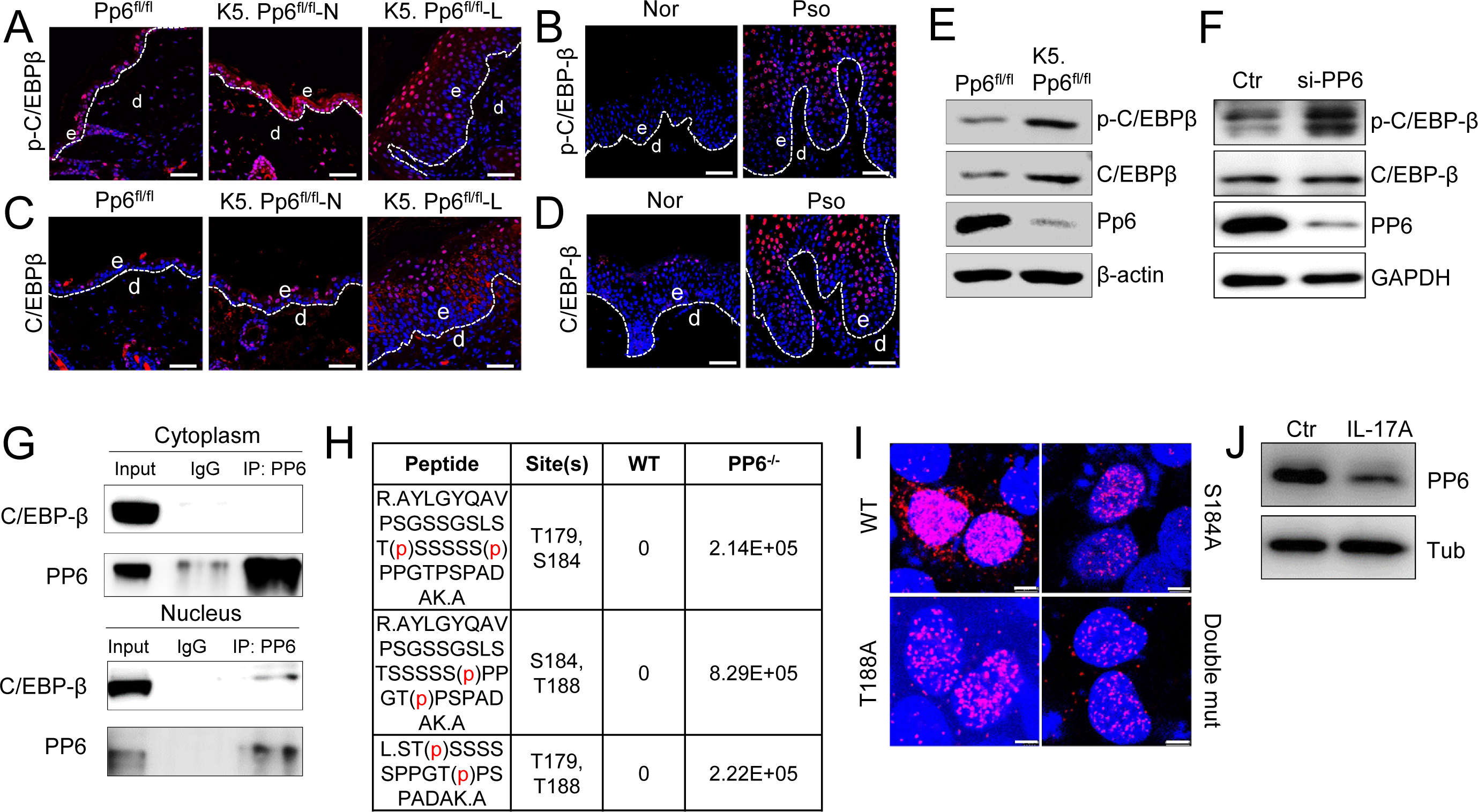
Loss of PP6 in keratinocytes activates C/EBP-β. (**A** and **B**) Immunofluorescent labeling of p-C/EBP-β in mouse (**A**) and human (**B**) skin sections. (**C** and **D**) Immunofluorescent labeling of C/EBPβ in mouse (**C**) and human (**D**) skin sections. (**E**) Immunoblot analysis of p-C/EBPβ, C/EBPβ and Pp6 expression in primary murine keratinocytes derived from Pp6^fl/fl^ and K5. Pp6^fl/fl^ mice cultured *in vitro* for 3 days. (**F**) Immunoblot analysis of p-C/EBP-β, C/EBP-β and PP6 expression in NHEKs transfected with a scrambled siRNA (Ctr) or with PP6 siRNA (si-PP6) for 48 h. (**G**) Nuclear and cytoplasmic extracts from HaCaT keratinocytes were immunoprecipitated with an anti-PP6 antibody. The immunoprecipitates were immunoblotted with anti-PP6 and anti-C/EBP-β. (**H**) LC-MS/MS-based detection of phosphorylation sites on C/EBP-β immunoprecipitated from WT 293FT cells and PP6^-/-^ 293FT cells. (**I**) Confocal visualization of PLA signals in HaCaT cells over-expressing WT C/EBP-β, C/EBP-β^S184A^, C/EBP-β^T188A^ or C/EBP-β^S184A^, plus C/EBP-β^T188A^ (double mut). (**J**) Immunoblot analysis of PP6 expression in NHEKs stimulated with recombinant IL-17A for 12 h or un-stimulated (Ctr). For (**A**-**D**), the dotted lines indicate the border between the epidermis and the dermis; Pp6^fl/fl^, healthy skin sample from control mice; K5. Pp6^fl/fl^-N, unaffected skin from K5. Pp6^fl/fl^ mice; K5. Pp6^fl/fl^-L, affected skin from K5. Pp6^fl/fl^ mice; Nor, normal human skin; Pso, psoriatic human skin; the scale bars represent 50 μm in (**A**-**D**) and 5 μm in (**I**). The data (**A**-**D**) are representative of > 5 samples in each group. The data (**E**-**G**, **I** and **J**) are representative of three independent experiments.

**Figure S4.**
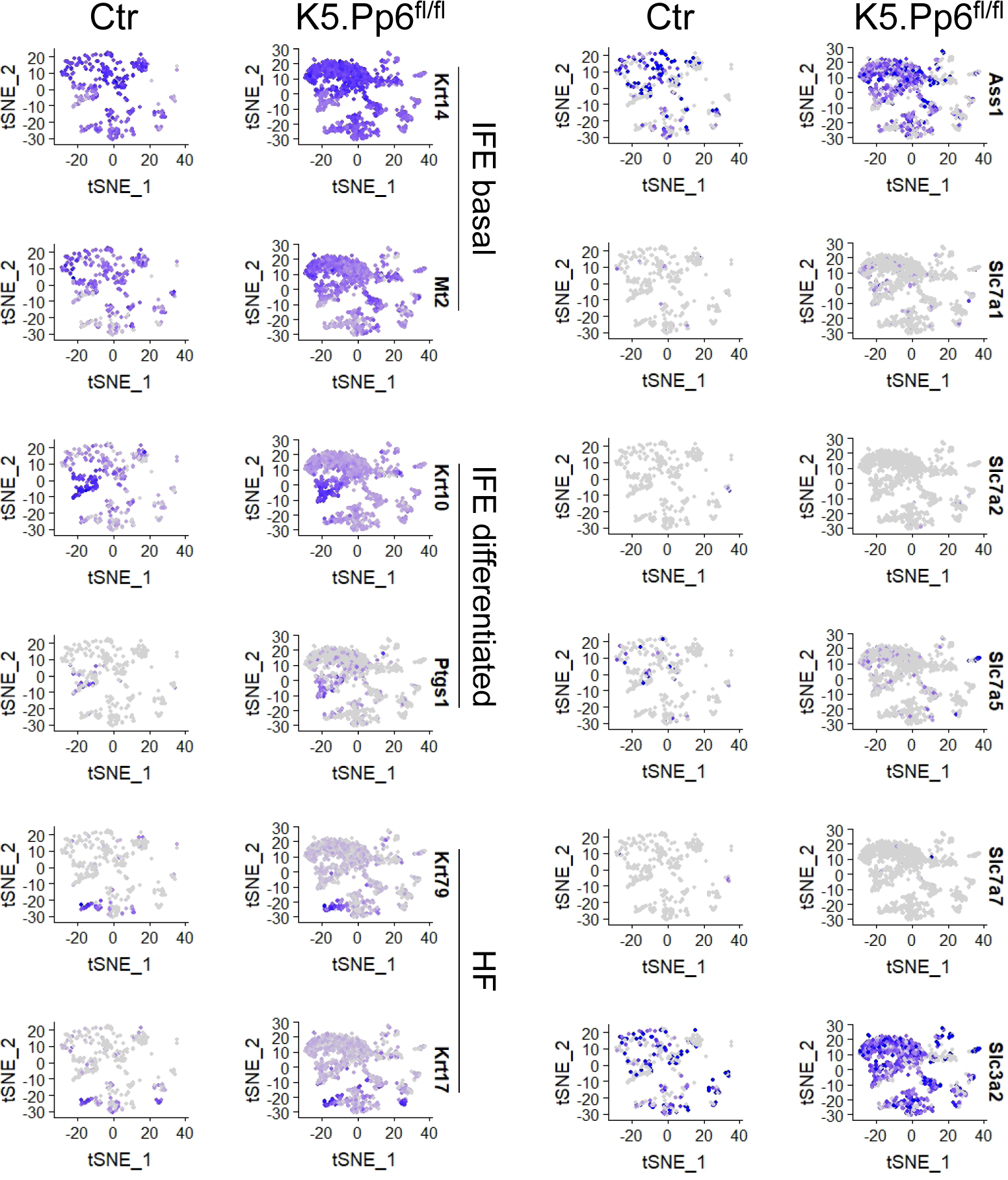
Mouse IFE basal keratinocytes uniquely and highly express Ass1. Feature-plots of the indicated genes in merged scRNAseq datasets of epidermal cells from C57BL/6 mice and K5. Pp6^fl/fl^ mice. IFE, interfollicular epidermis; HF, hair follicle.

**Figure S5.**
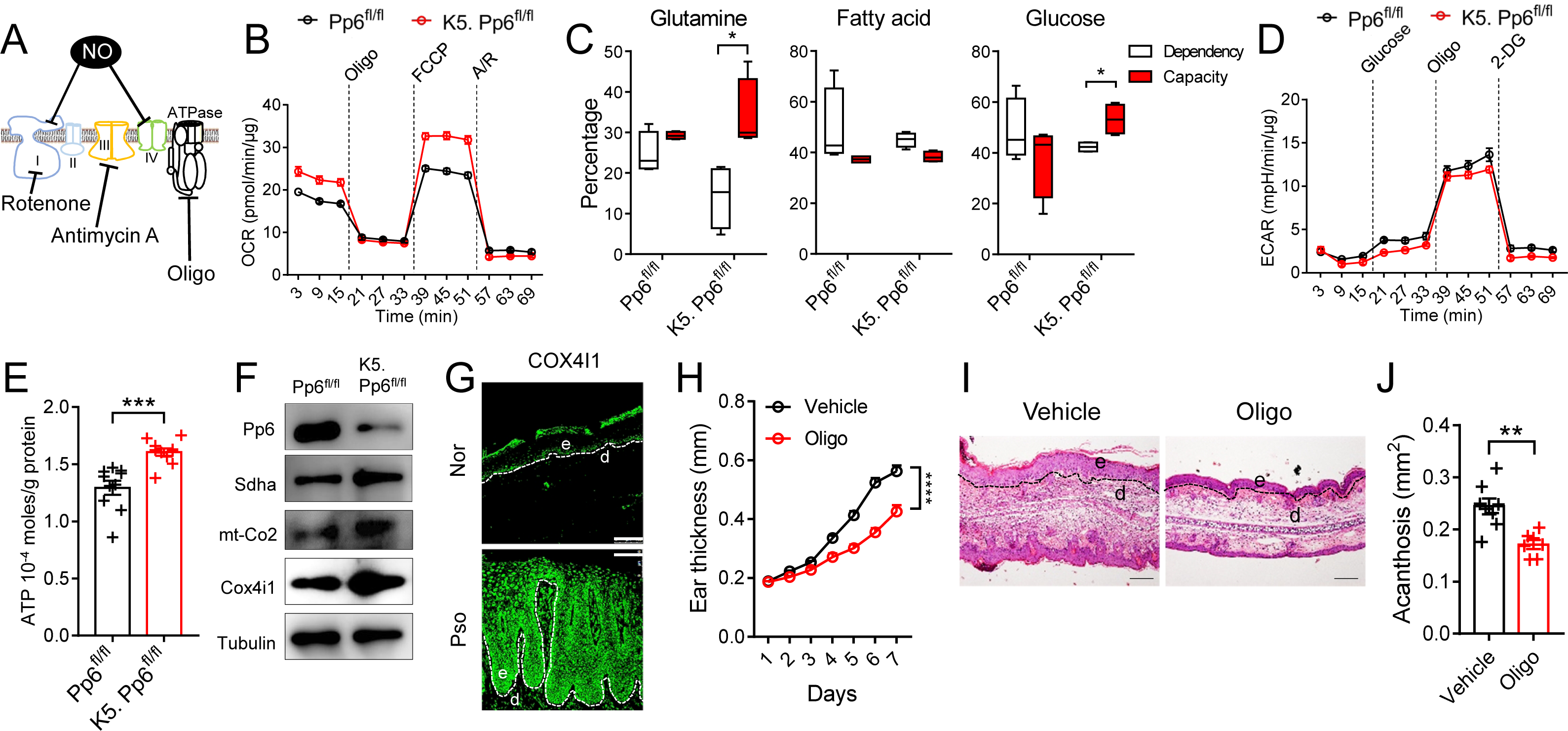
Pp6-deficiency keratinocytes show enhanced OXPHOS capacity. (**A**) The inhibitory molecules that target ETC components. (**B**) OCR of Pp6^fl/fl^ and K5. Pp6^fl/fl^ keratinocytes sequentially treated with the indicated reagents (n = 5-6). (**C**) Mitochondrial fuel dependency and capacity of Pp6^fl/fl^ and K5. Pp6^fl/fl^ keratinocytes. The difference between the dependency and capacity for each fuel represents flexibility (n = 4). (**D**) ECAR of Pp6^fl/fl^ and K5. Pp6^fl/fl^ keratinocytes sequentially treated with indicated reagents (n = 5-6). (**E**) Cellular ATP levels in Pp6^fl/fl^ and K5. Pp6^fl/fl^ keratinocyte measured by luminescence-based quantification (n = 10). (**F**) Immunoblot analysis of ETC components in Pp6^fl/fl^ and K5. Pp6^fl/fl^ keratinocytes. (**G**) Immunofluorescent labeling of COX4I1 in human skin sections. The data are representative of > 5 samples in each group. (**H**) Changes in ear thickness in IMQ-induced C57BL/6 mice treated with vehicle or Oligomycin (n = 8). (**I**) Representative H&E staining of mouse ears from IMQ-induced C57BL/6 mice treated with vehicle or Oligomycin. (**J**) Acanthosis of mouse ears from IMQ-induced C57BL/6 mice treated with vehicle or Oligomycin. For (**G** and **I**), the dotted lines indicate the border between the epidermis and the dermis; Nor, normal human skin; Pso, psoriatic human skin; scale bars represent 100 μm. Oligo, Oligomycin; FCCP, Trifluoromethoxy carbonyl cyanide phenylhydrazone; A/R, Antimycin and Rotenone; 2-DG, 2-Deoxy-D-glucose. The data (**B**, **D** and **E**) are normalized to the total protein content of each sample. The data (**B-J**) are representative of three independent experiments. \**p* < 0.05, \*\**p* < 0.01, \*\*\**p* < 0.001, \*\*\*\**p* < 0.0001, two-way ANOVA (**h**) and two-tailed Student’s *t*-test (means ± SEM).

**Figure S6.**
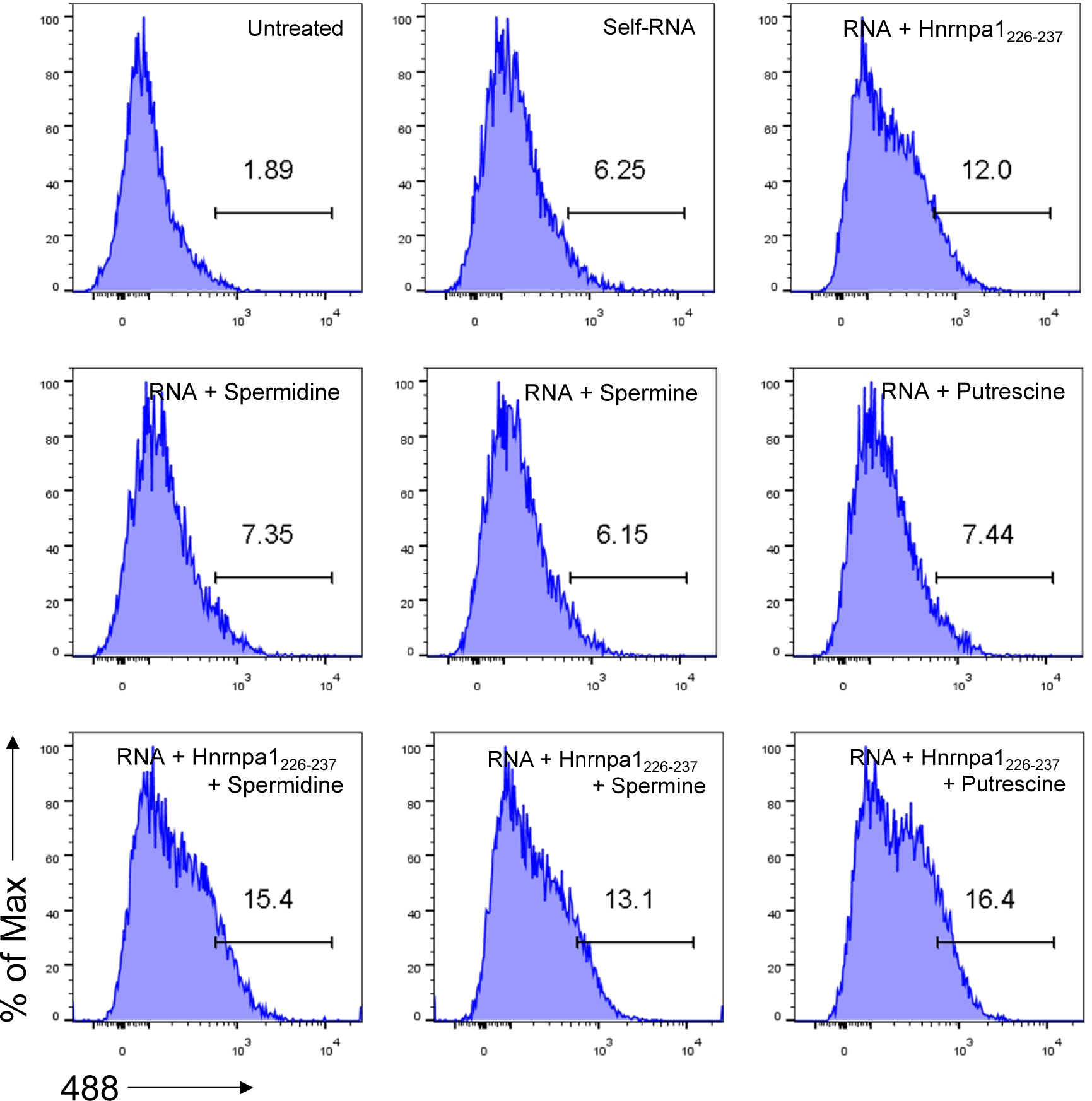
Self-RNA-polyamine-Hnrnpa1226-237 complexes are internalized by BMDCs. Flow cytometric analysis of BMDCs treated with 488-labeled RNA in complex with the indicated molecules. The data are representative of two independent experiments.

**Figure S7.**
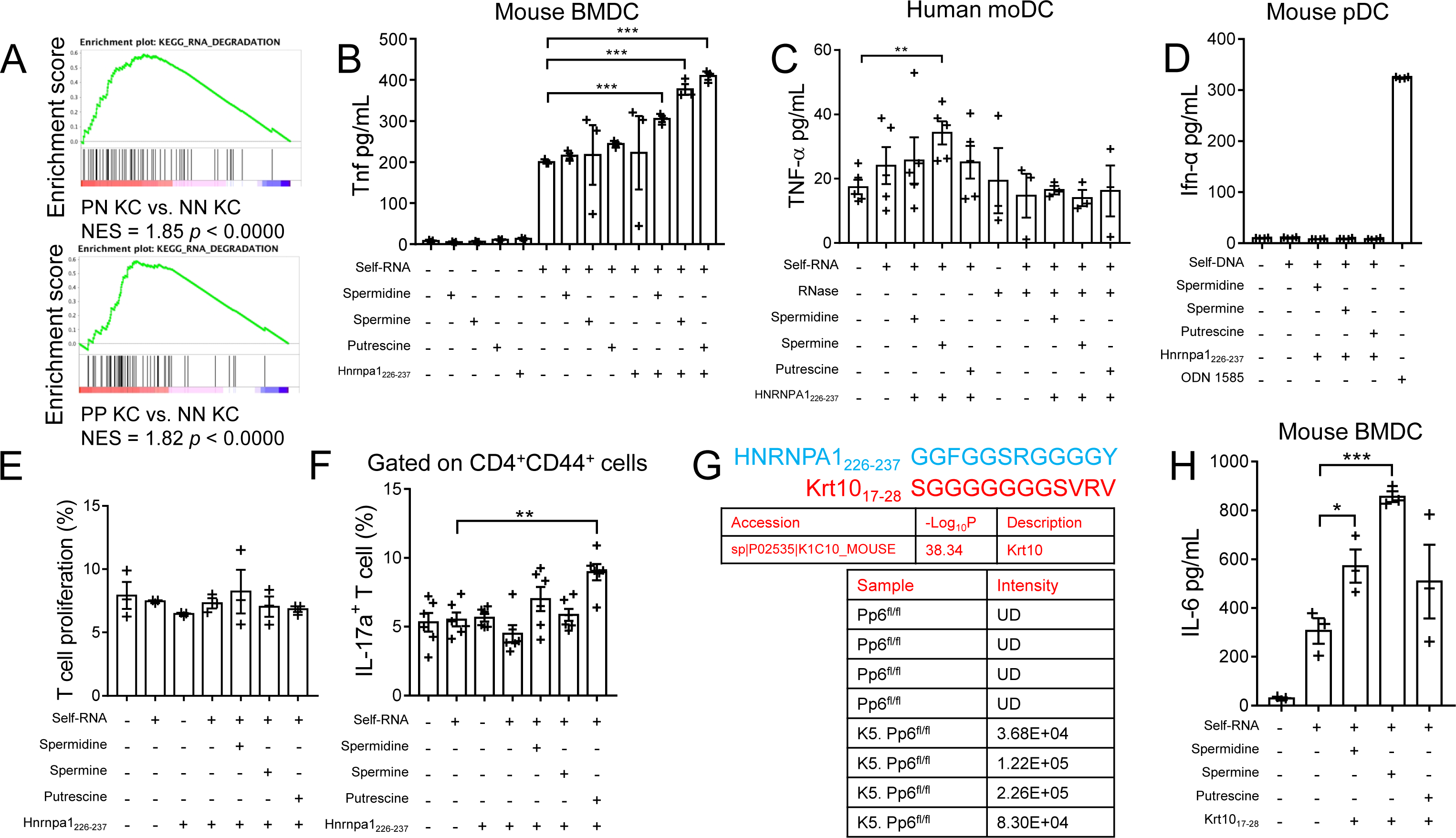
Self-RNA-polyamine-cargo peptide complexes activate BMDCs and induce subsequent T17 cell polarization. (**A**) GSEA of all the genes sequenced from cultured human keratinocytes (GSE107871) derived from healthy skin (NN KC), uninvolved skin of patients with psoriasis (PN KC) and involved skin of patients with psoriasis (PP KC) with genes involved in the RNA degradation pathway enriched. (**B**) ELISA quantification of Tnf in mouse BMDC cultures stimulated with the indicated reagents (n = 3). (**C**) ELISA quantification of TNF-α in human moDC cultures stimulated with the indicated reagents (n =3∼5). (**D**) ELISA quantification of Ifn-α in mouse pDC cultures stimulated with the indicated reagents (n = 4). (**E**) Statistics of T-cell proliferation rates (indicated by CD4^+^CD44^+^Violet^-^ cell percentages) in co-culture with BMDCs pre-treated with the indicated reagents (n = 3). (**F**) Statistics of IL-17a^+^ cell percentages in co-culture with BMDCs pre-treated with the indicated reagents (n = 6). (**G**) LC-MS/MS-based detection of Krt1017-28 abundance in the sera of Pp6^fl/fl^ mice (n = 4) and K5. Pp6^fl/fl^ mice (n = 4); UD, undetected. (**H**) ELISA quantification of IL-6 in mouse BMDC cultures stimulated with the indicated reagents (n = 3). The data (**B**-**F** and **H**) are representative of three independent experiments. \**p* < 0.05, \*\**p* < 0.01, \*\*\**p* < 0.001, two-tailed Student’s *t*-test (means ± SEM).

**Figure S8.**
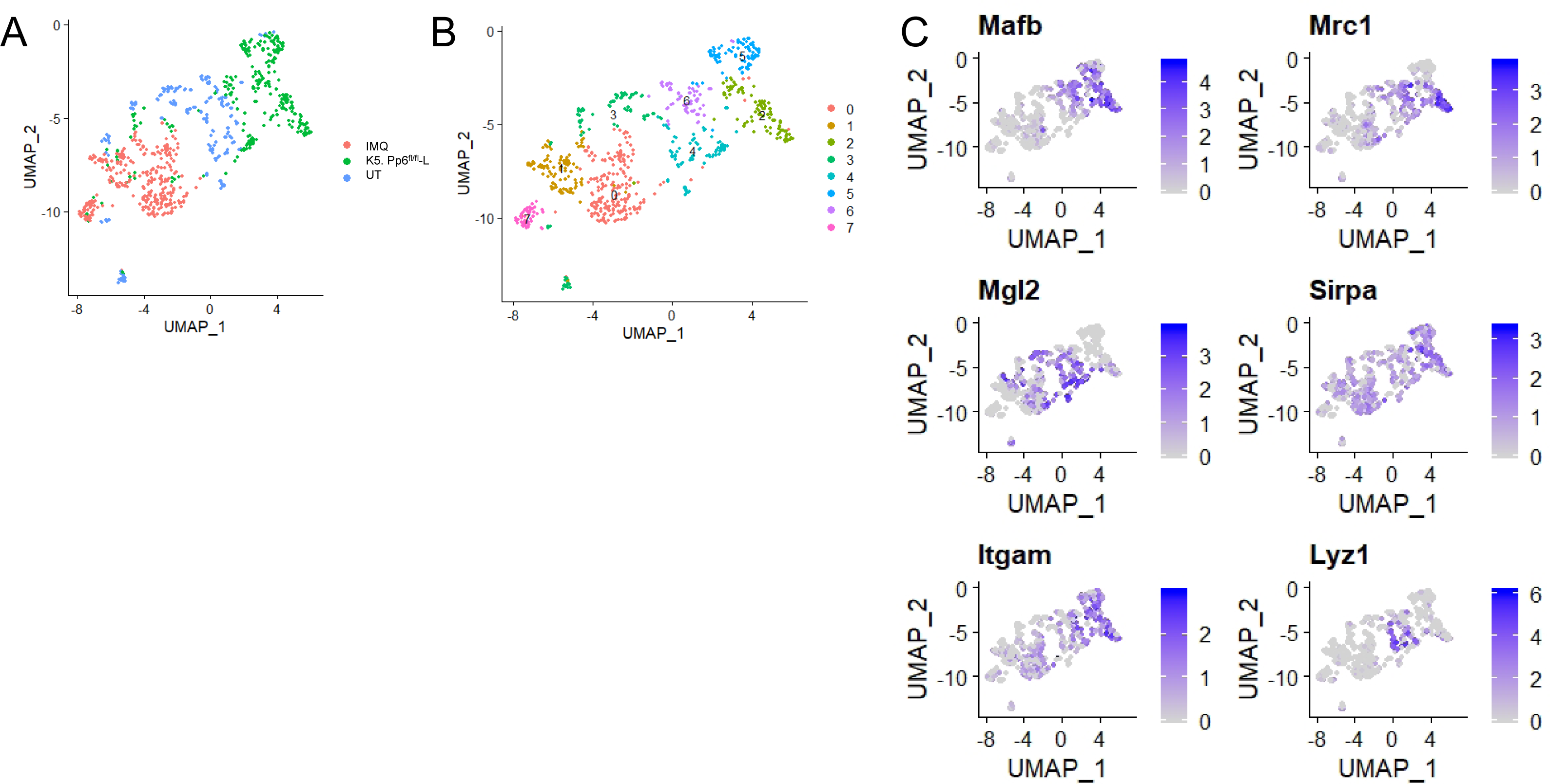
Dominant DC populations in diseased K5. Pp6^fl/fl^ skin and IMQ-treated mouse skin. (**A**-**B**) t-SNE plots of merged scRNAseq datasets of skin DCs derived from diseased K5. Pp6^fl/fl^ mice, untreated C57BL/6 mice (UT) and C57BL/6 mice treated with IMQ (IMQ), colored separately by samples (**A**) or cell sub-clusters (**B**). (**C**) Feature-plots of the indicated genes in merged scRNAseq datasets of skin DCs derived from diseased K5. Pp6^fl/fl^ mice, UT C57BL/6 mice and IMQ C57BL/6 mice.

**Figure S9.**
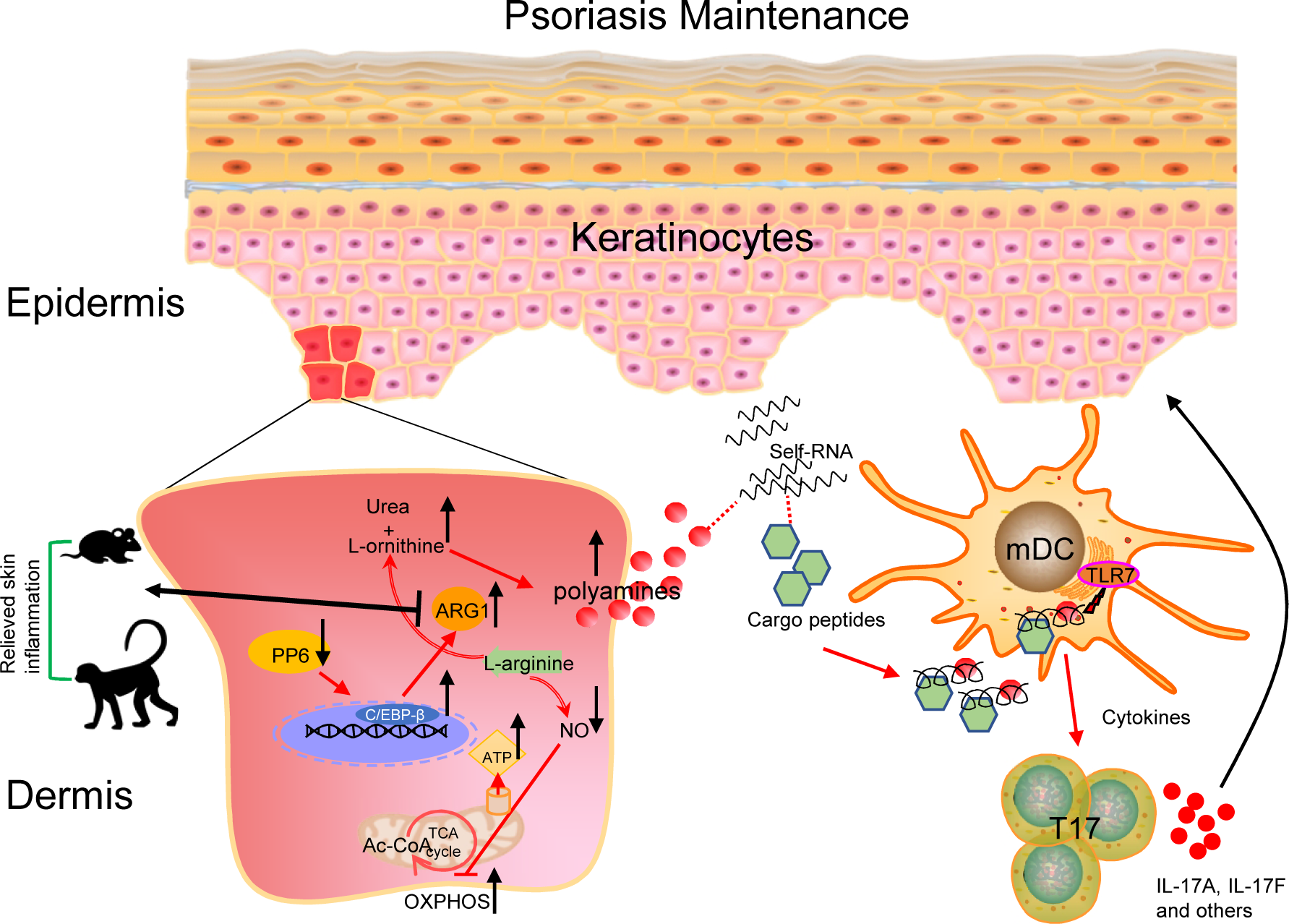
Summary of the study. Hyperproliferating keratinocytes upregulate ARG1 and undergo urea cycle rewiring, leading to local polyamine accumulation. In complex with HNRNPA1226-237 identified in the blood plasma of patients with psoriasis, polyamines aggregate self-RNA that is released by psoriatic keratinocytes, facilitating self-RNA endosomal sensing by myeloid dendritic cells and initiating downstream inflammatory cascades. Inhibiting ARG1, a critical regulator of the urea cycle, markedly improves psoriasis-like skin inflammation in mice and non-human primates.

